# Comparative transcriptome of normal and cancer-associated fibroblasts

**DOI:** 10.1101/2024.03.18.585496

**Authors:** Apoorva Abikar, Mohammad Mehaboob Subhani Mustafa, Radhika Rajiv Athalye, Namratha Nadig, Ninad Tamboli, Vinod Babu, Ramaiah Keshavamurthy, Prathibha Ranganathan

## Abstract

The characteristics of a tumor are largely determined by its interaction with the surrounding micro-environment (TME). TME consists of both cellular and non-cellular components. Cancer associated fibroblasts (CAFs) are a major component of the TME. They are a source of many secreted factors that influence the survival and progression of tumors as well as their response to drugs. Identification of markers either overexpressed in CAFs or unique to CAFs would pave the way for novel therapeutic strategies that would combine conventional chemotherapy and TME-targeted therapy for a better outcome. We have used fibroblast derived from Benign Prostatic Hyperplasia (BPH) and prostate cancer to perform a transcriptome analysis in order to get a comparative profile of normal and cancer-associated fibroblasts. This has identified 818 differentially expressed mRNAs and 17 lincRNAs between normal and cancer-associated fibroblasts.

## Introduction

The tumor microenvironment (TME) is a complex and dynamic entity, which is shaped by the interactions with the tumor itself. Although the framework and composition of the TME may vary according to the tumor type, some hallmarks of the TME remain the same. The TME is comprised of both cellular and non-cellular components, such as growth factors, cytokines, ECM (extracellular matrix) proteins, and metabolites [1]. The dynamic and reciprocal interactions between the tumor and its microenvironment influence cancer cell survival, local invasion, metastatic spread [2, 3], immune surveillance, angiogenesis [4] as well as response to therapy [5, 6].

One of the major cell types in the TME is cancer-associated fibroblasts. CAFs are a major source of growth factors, cytokines, and other signaling molecules, which impact cancer cell behavior [6]. When subjected to chemotherapy, along with the cancer cells, the CAFs also are subjected to changes. These therapies are likely to stimulate CAFs to release factors that could influence the stemness, metabolic status, signaling cascades, etc. within the tumor which can prevent cancer cell eradication and perhaps cause recurrence [7].

CAFs are pivotal in driving cancer progression through their involvement in processes such as extracellular matrix (ECM) deposition and remodeling, extensive communication with cancer cells, promoting epithelial-to-mesenchymal transition (EMT), facilitating invasion, metastasis, and even contributing to therapy resistance [8]. CAFs are also recognized for their involvement in developing resistance to anti-cancer therapy by providing a protective environment for tumor cells. There exists a symbiotic relationship between tumor cells and CAFs, wherein CAFs provide the necessary resources for tumor cell growth and survival, thereby contributing to the development of a chemoresistant phenotype [9]. Considering the pleiotropic effects of the tumor microenvironment, particularly CAFs, an insight into the specific factors responsible for therapeutic resistance can potentially pave the way for newer and more effective strategies for treatment. The major hurdle in this direction is the lack of distinguishing biomarkers for CAFs that would allow for their exclusive targeting. There is a high heterogeneity of CAF functions-both pro-tumorigenic and anti-tumorigenic within the same tumor [10]. Hence targeting the CAFs/derived factors has to be done with extreme caution to avoid adverse effects. Additionally, more than just targeting CAFs might lead to significant clinical benefits, as pro-tumorigenic CAFs can support tumor progression, but they may not be indispensable for tumor growth and survival. In other words, tumor cells may not solely depend on the presence of CAFs. CAF-targeted therapy in combination with other chemotherapeutic drugs is likely to improve treatment outcomes. In this study, we have used transcriptome analysis to identify differentially expressed genes and non-coding RNAs between normal and cancer associated fibroblasts on a human prostate model.

## Materials and Methods

### Clinical specimen collection

All patient samples were collected from the Institute of Nephro-Urology (Department of Urology), Bengaluru. The study was approved by the Institutional Ethics committee of both the institutions and informed consent has been taken from all the participants. Samples were taken either by TRUS (Transrectal Ultrasound scan) or TRUP (Transurethral resection of the prostate) methods. Classification of the samples as benign or malignant was done by pathologists as per standard criteria.

### Culturing of Fibroblasts

Surgical or biopsy specimens were rinsed thoroughly in sterile saline and transferred to transport media (RPMI 1640 (Cat No: 23400-021) with 2X PenStrep (Cat No: 15140122)). Subsequently, these specimens were rinsed thoroughly with RPMI media containing antibiotics, minced into fine pieces and transferred to culture flasks keeping sufficient distance between each piece for the cells to migrate out. Media was changed periodically. Once the fibroblasts migrated out of the tissues, cells were transferred to fresh flasks.

### RNA isolation

Total RNA was isolated from the cultured fibroblasts using a Qiagen RNeasy kit (Cat No: 74104) according to the manufacturer’s instructions. RNA was quantified on nanodrop (Thermo Scientific™ NanoDrop™ One Microvolume UV-Vis spectrophotometer). The quality of RNA samples was assessed by running them on 1% agarose gel.

### RNA sequencing

RNA sequencing was outsourced to Wipro Life Science Labs, Bengaluru. Quality assessment was done using Agilent TapeStation and all samples had RIN >9. Samples were further taken for library preparation and RNA sequencing using the Illumina platform.

### RNA Sequencing Analysis Workflow

The quality assessment of the data was performed for base quality and contamination by sequencing artifacts. The adapters were trimmed and poor-quality sequences were filtered using Trim Galore. Trimmed sequence reads were mapped to reference genome (**Assembly**: hg38, GRCh38.p12 (GCA_000001405.27), Dec. 2017, **Data Source**: UCSC Genome Browser, Weblink:http://hgdownload.soe.ucsc.edu/goldenPath/hg38/bigZips/analysisSet/hg38.analysisSet.fa.gz) with splice aware alignment tool STAR. R subread R package was used to get feature-specific expression counts. Low-count features across the samples were detected and removed using the NOISeq R package followed by expression count normalization with the TMM method (from the NOISeq R package). Differential analysis was performed with the NOISeq R package where the group information was used to define biological replicates. Genes/transcripts (mRNA and lincRNA) were considered differentially expressed when they showed at least a log2 fold change of 1.5 differential expression between normal and CAFs.

### LINCRNA functional annotation

NPInter v5.0 (http://bigdata.ibp.ac.cn/npinter5/) [11, 12] is a database that provides collective information about the multidimensional interactions of ncRNAs (lincRNA, miRNA, circRNA, etc.) with protein, RNA, and DNA. This database contains information about RNA interactions based on literature mining and high-throughput sequencing data with functional annotation [11].

The list of differentially expressed lincRNAs from our experiment was fed into NPInter v5.0 database and segregated according to the interactors (proteins, mRNA, and ncRNA).

The miRNAs obtained from the lincRNA-miRNA (ncRNA) interaction (from NPInter v5.0 database) were further subjected to the miRDB database (https://mirdb.org/)[13] to predict its mRNA targets.

## Results

### Differential expression of genes between normal and cancer-associated fibroblasts

We have identified 818 genes and 17 long intergenic non-coding RNAs (lincRNAs) that exhibit differential expression between normal fibroblasts and CAFs with a minimum log2 fold of 1.5. Of these, 380 genes and 7 lincRNAs were found to be overexpressed (Table 1A), while 438 genes and 10 lincRNAs were under-expressed (Table 1B) in CAFs as compared to normal fibroblasts.

**Table 1:**
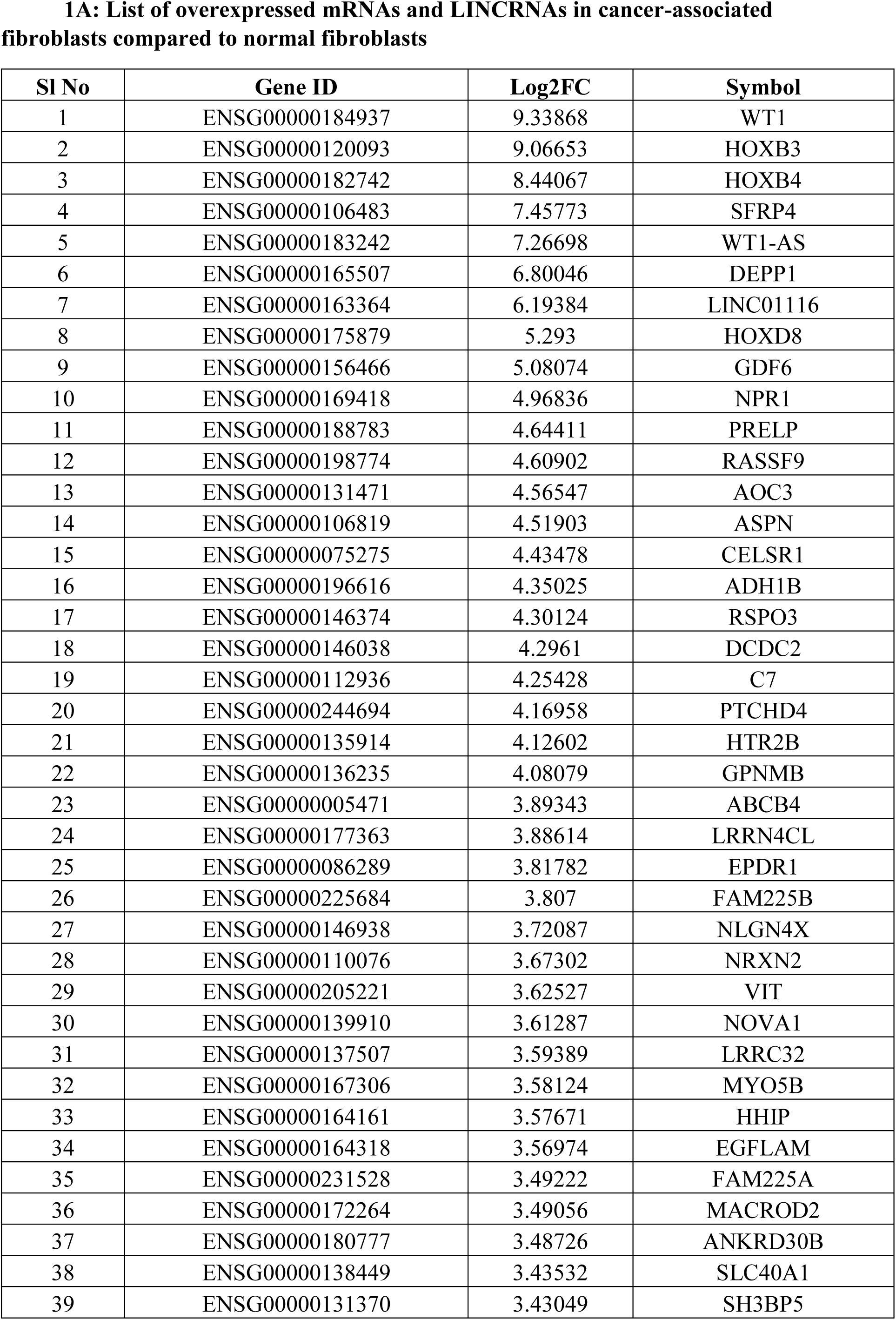
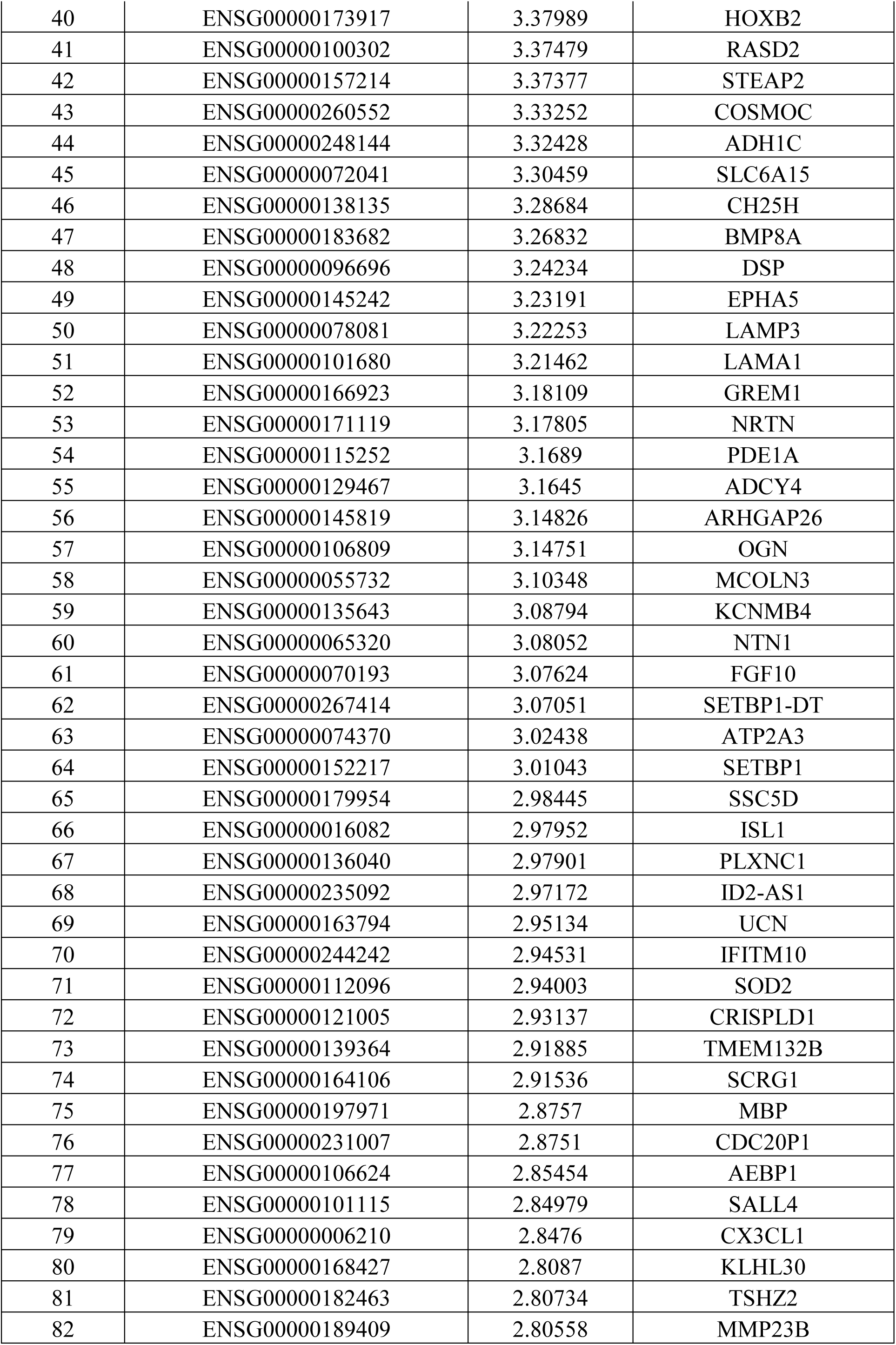
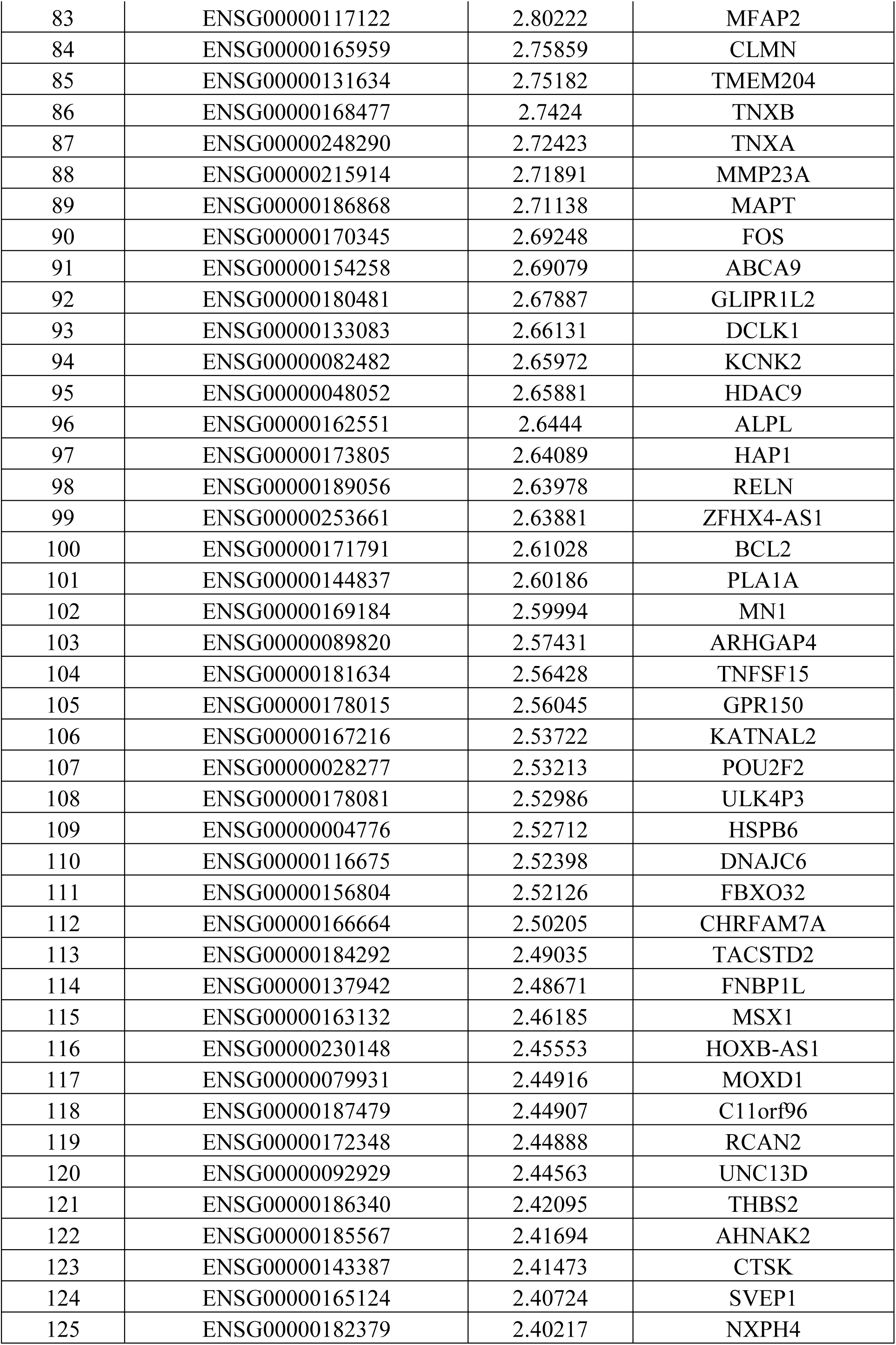
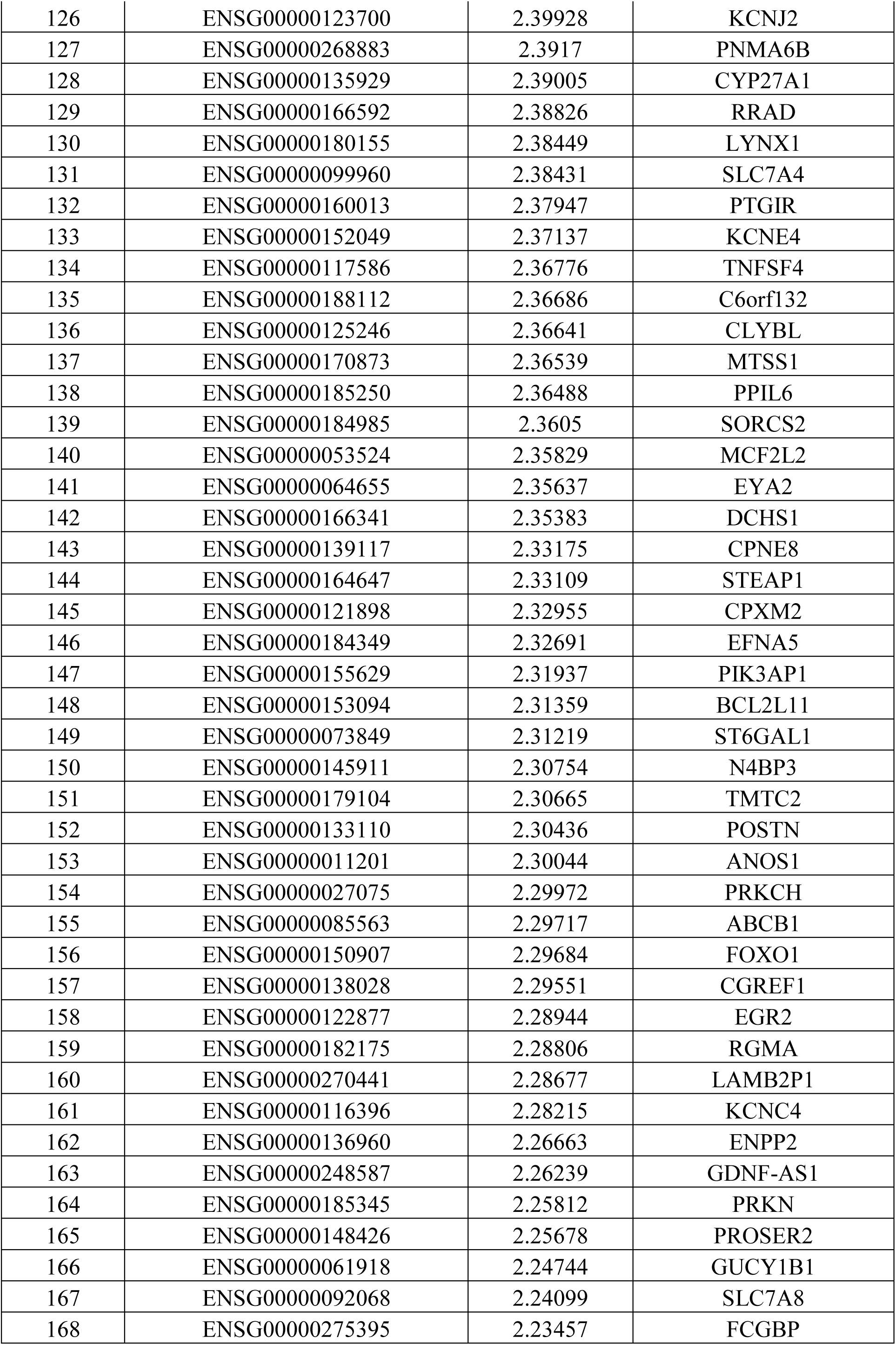
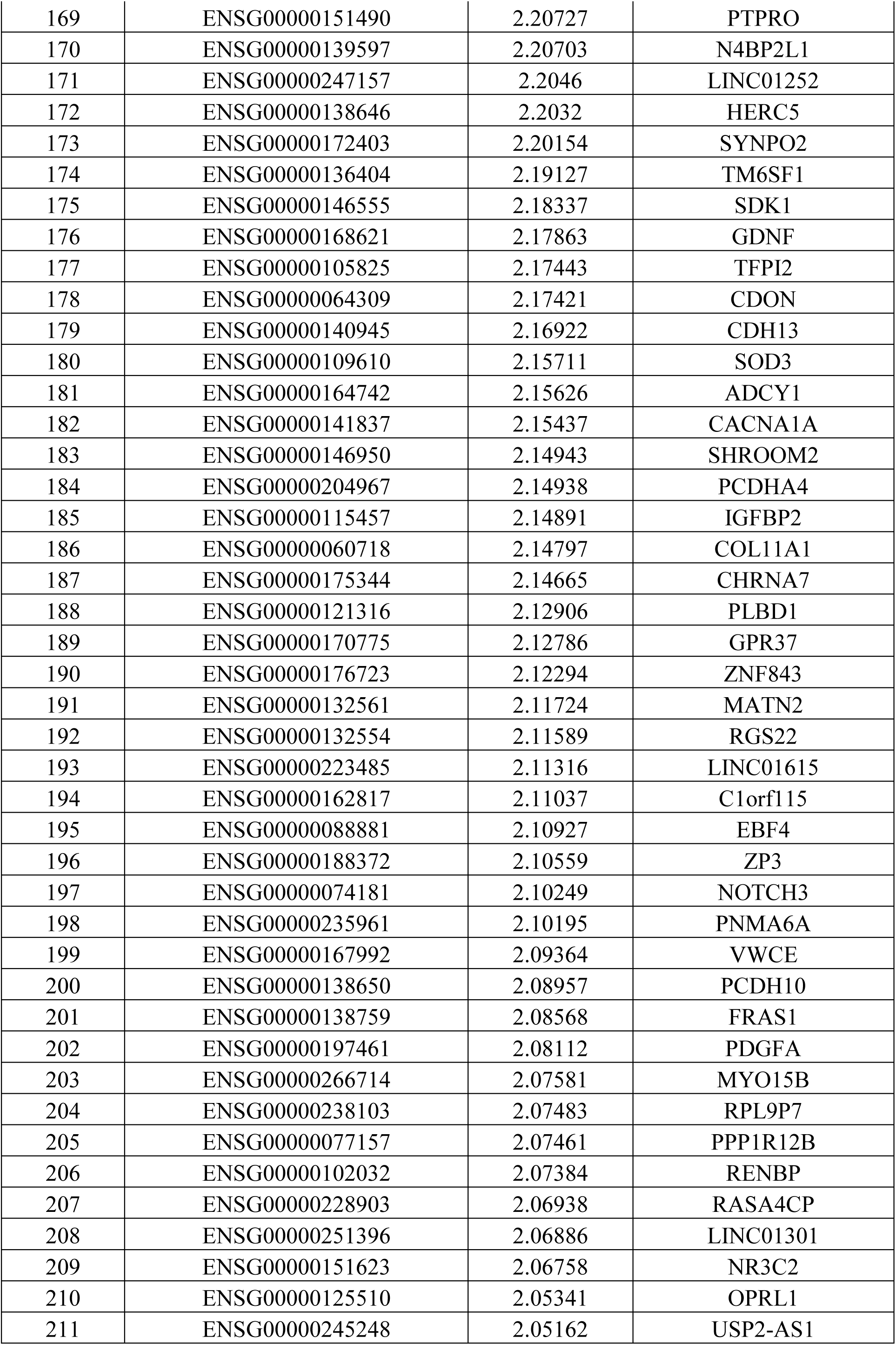
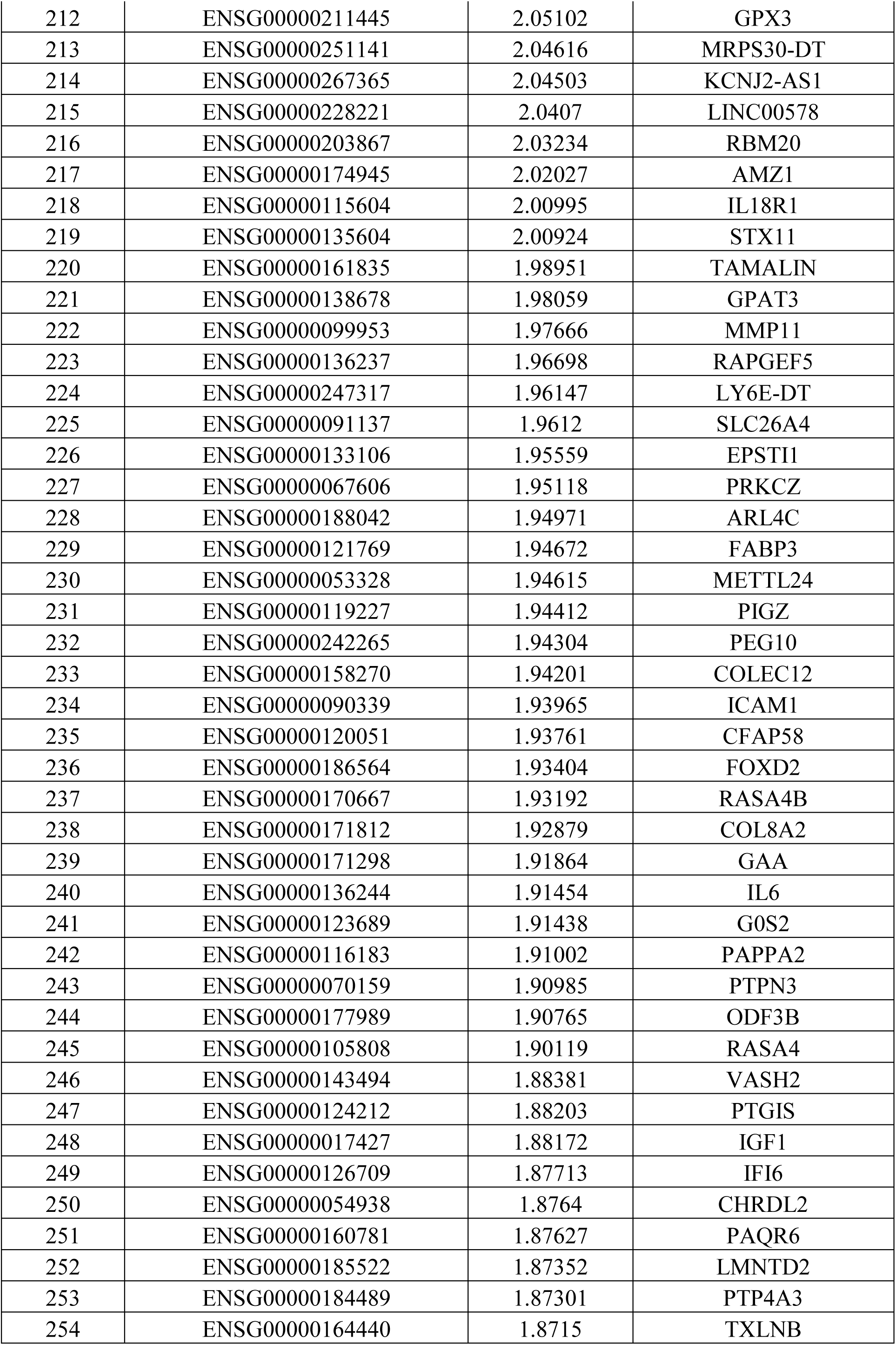
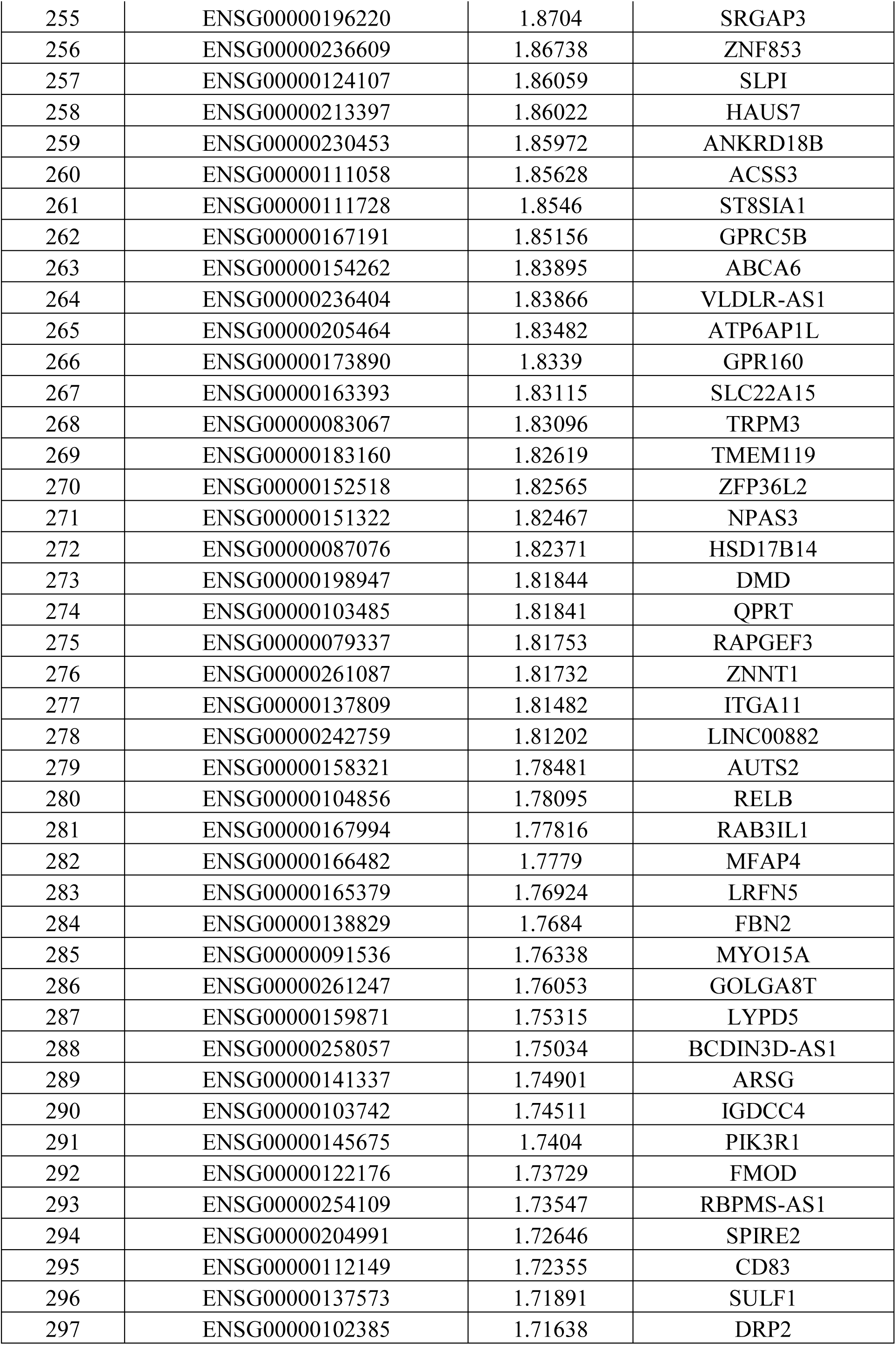
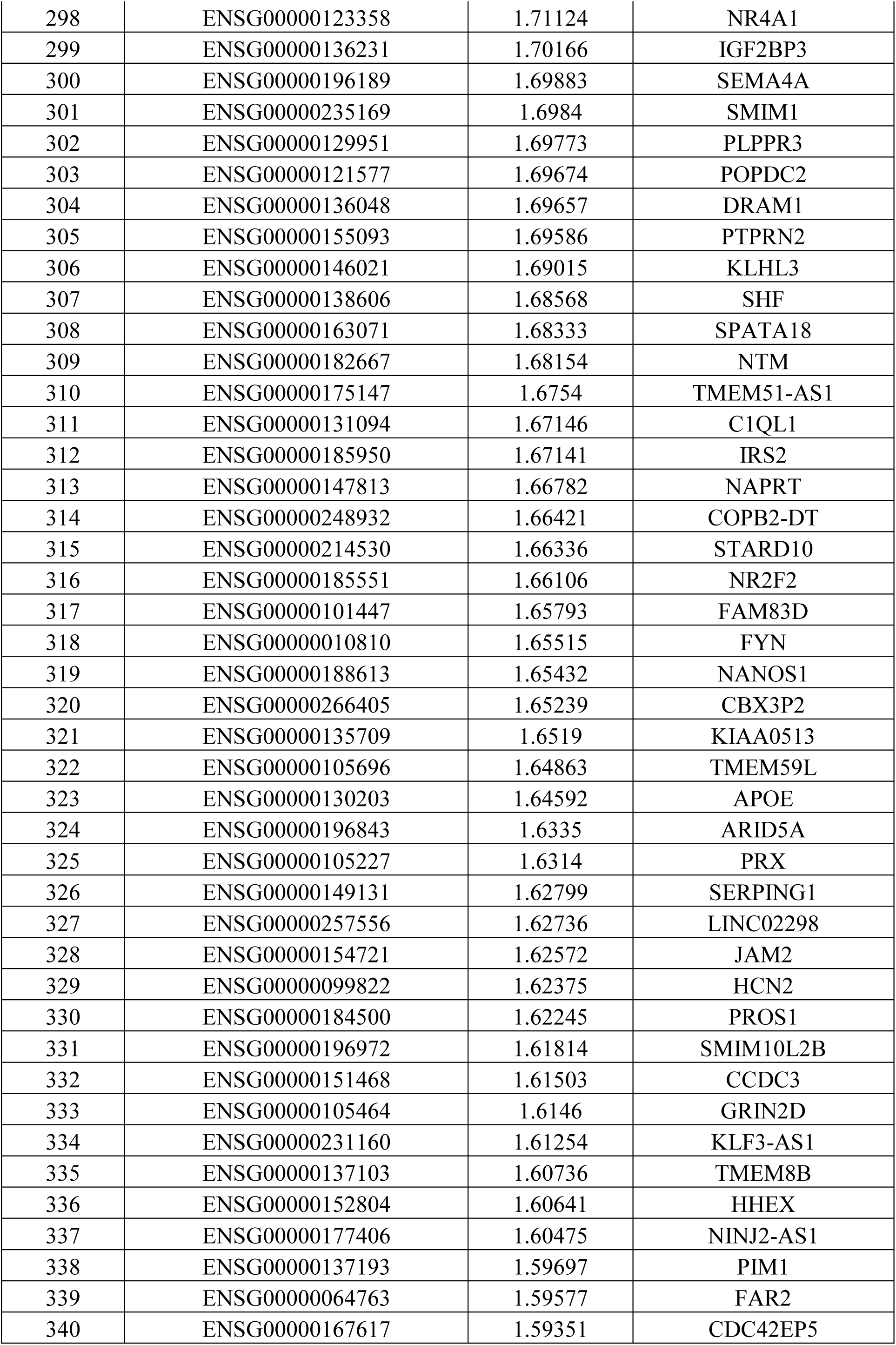
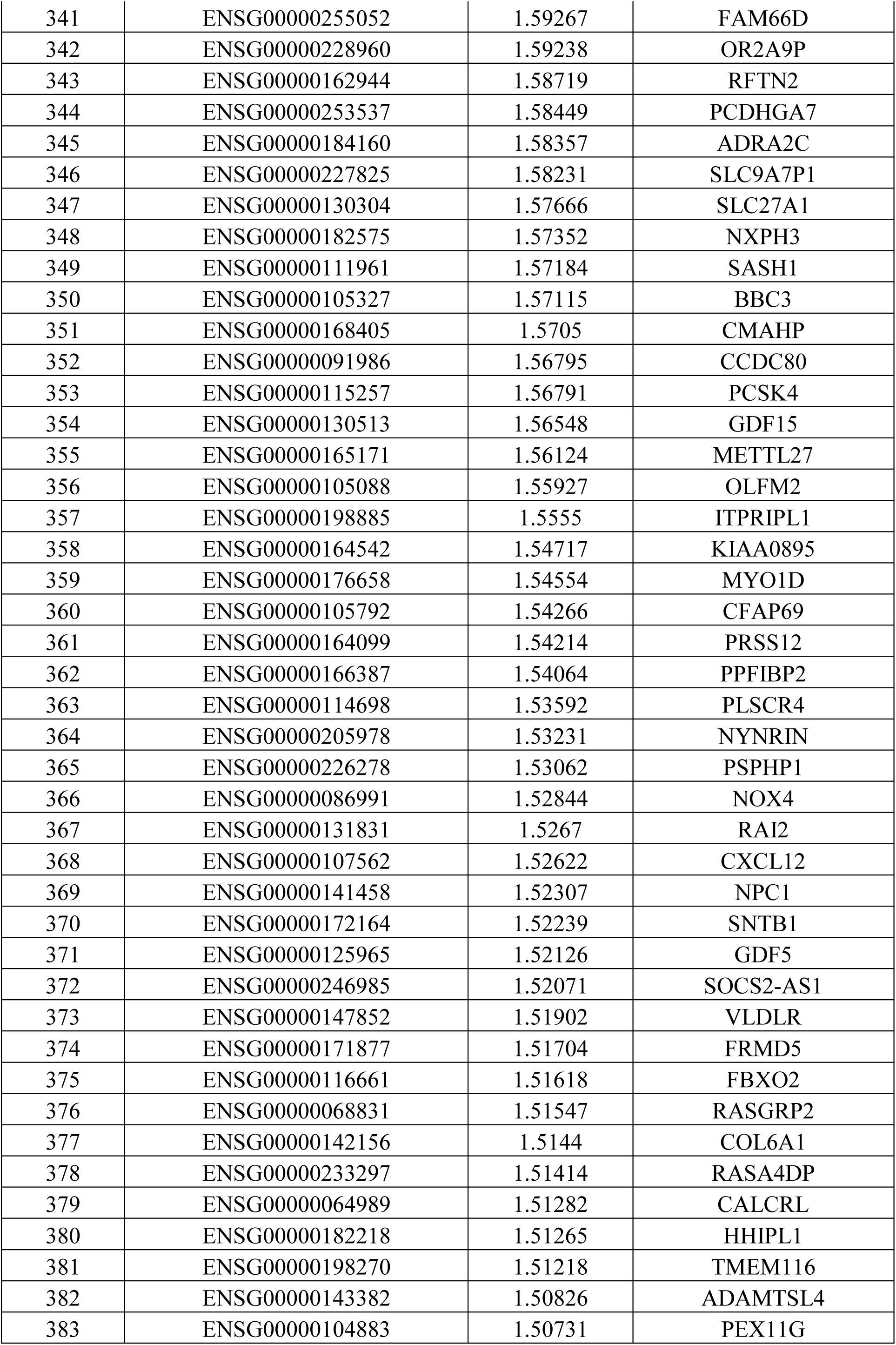
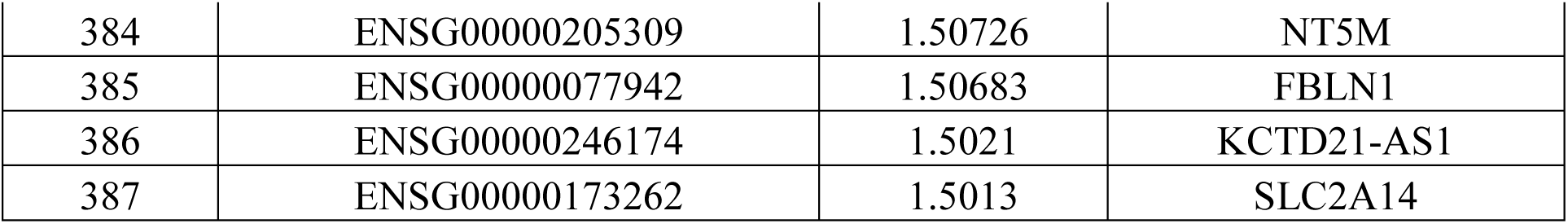
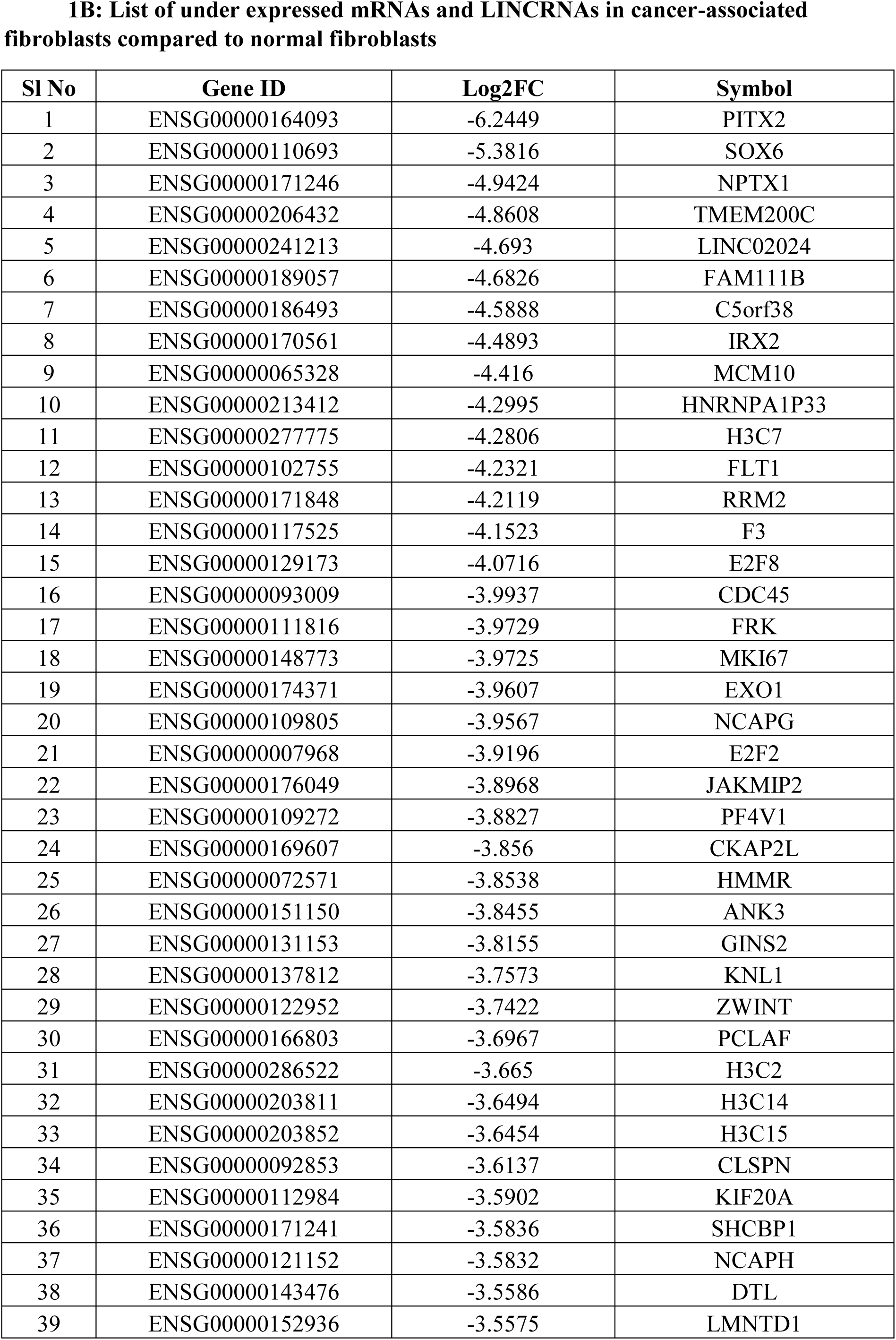
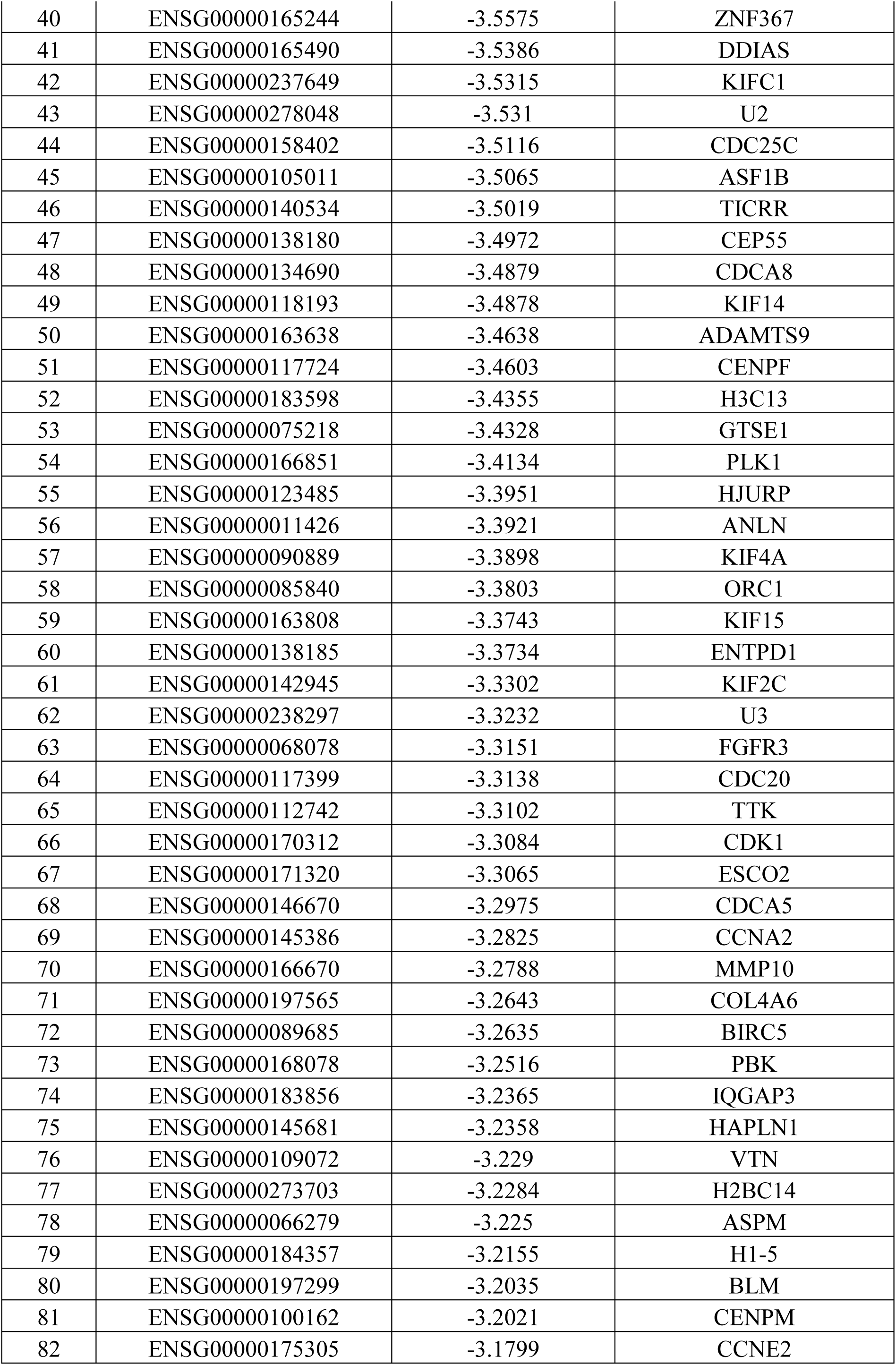
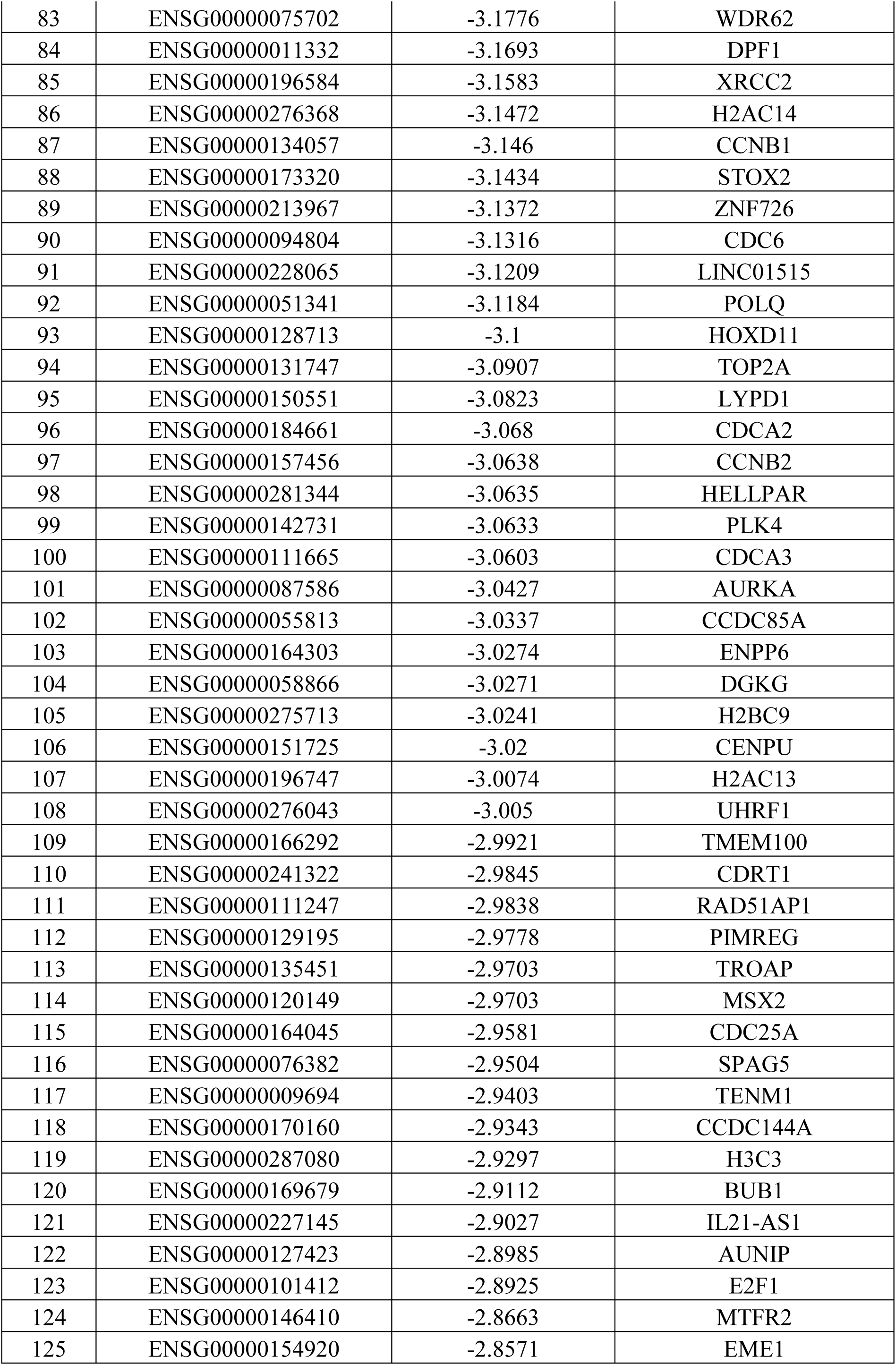
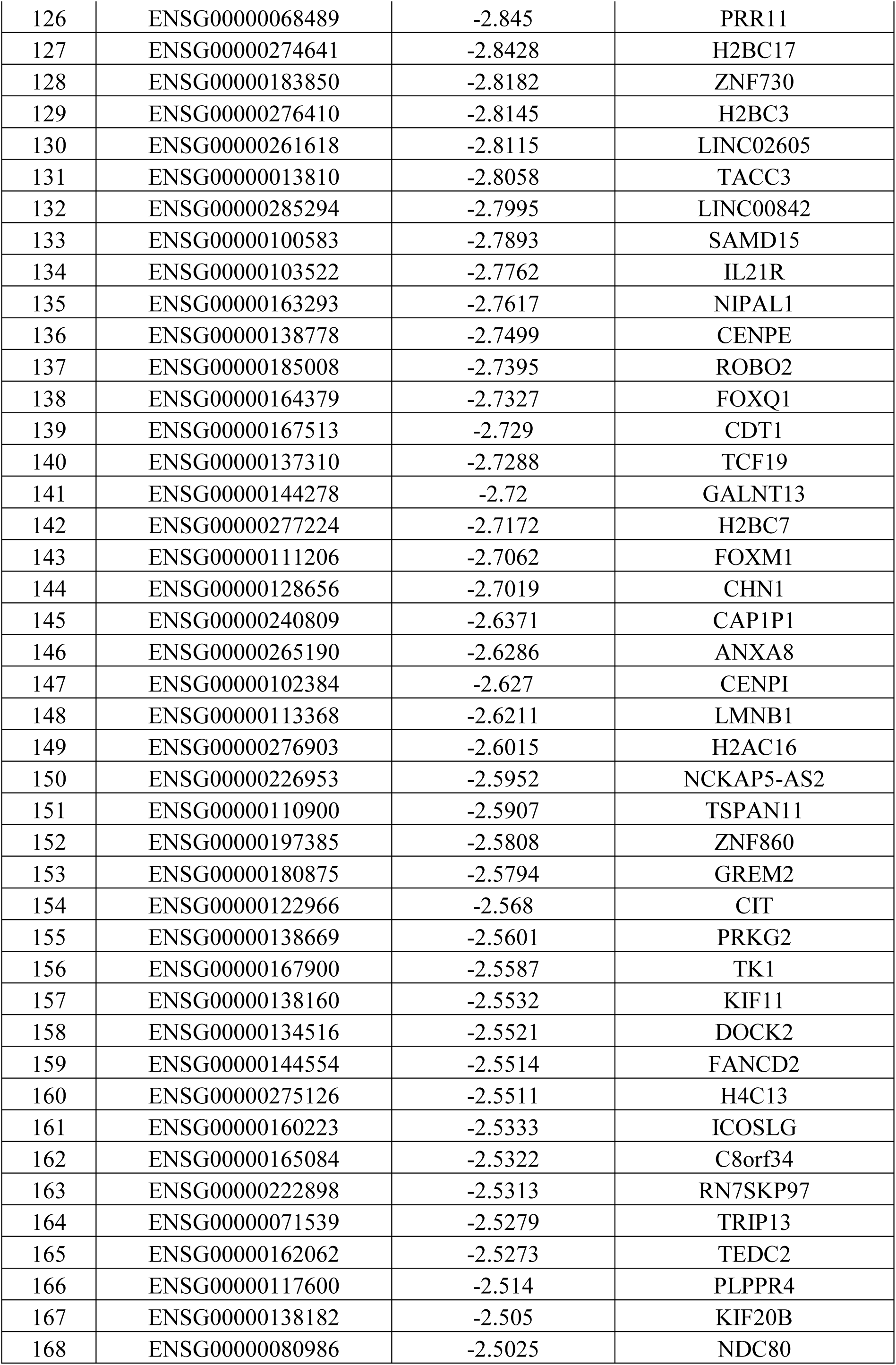
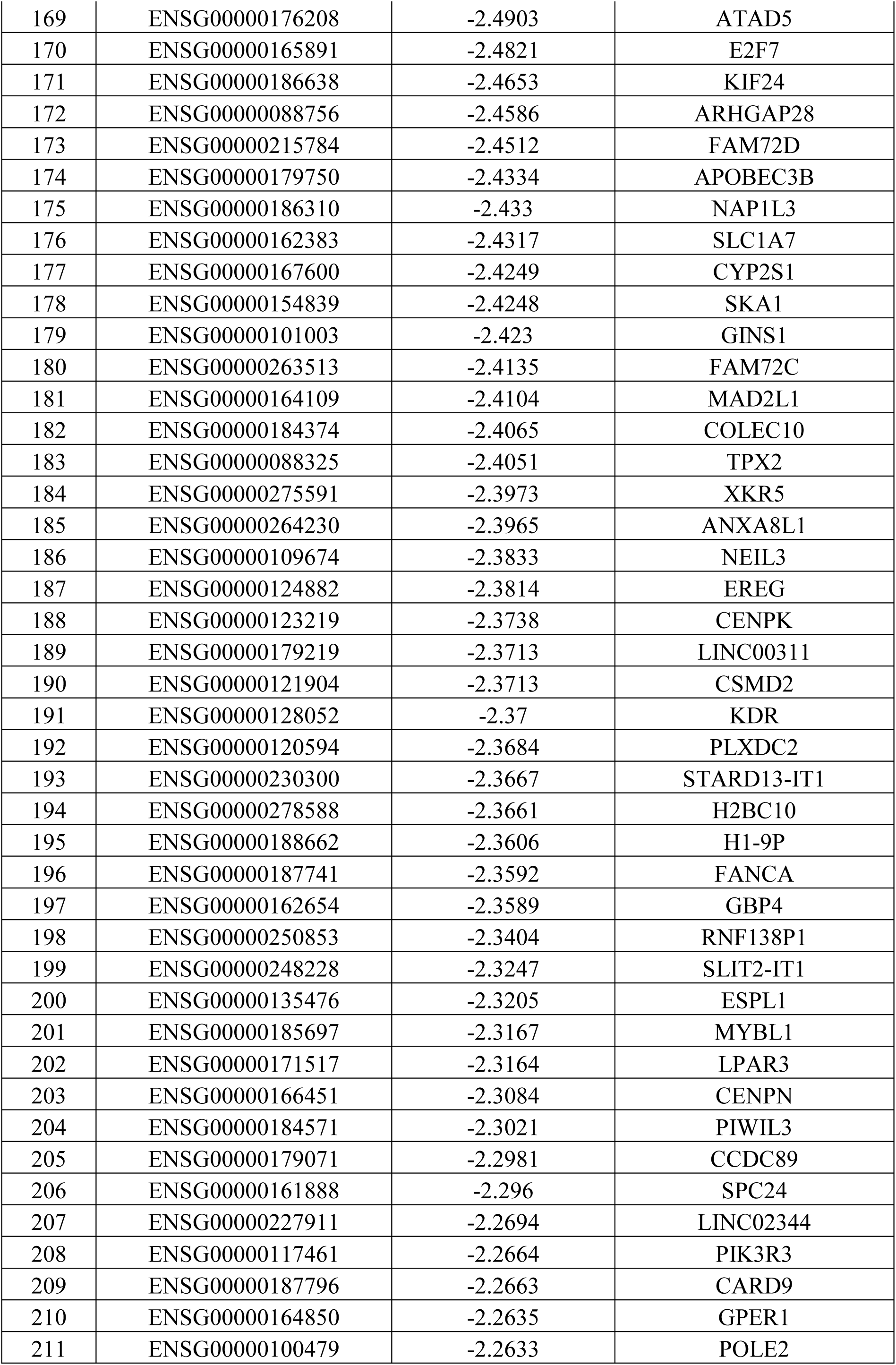
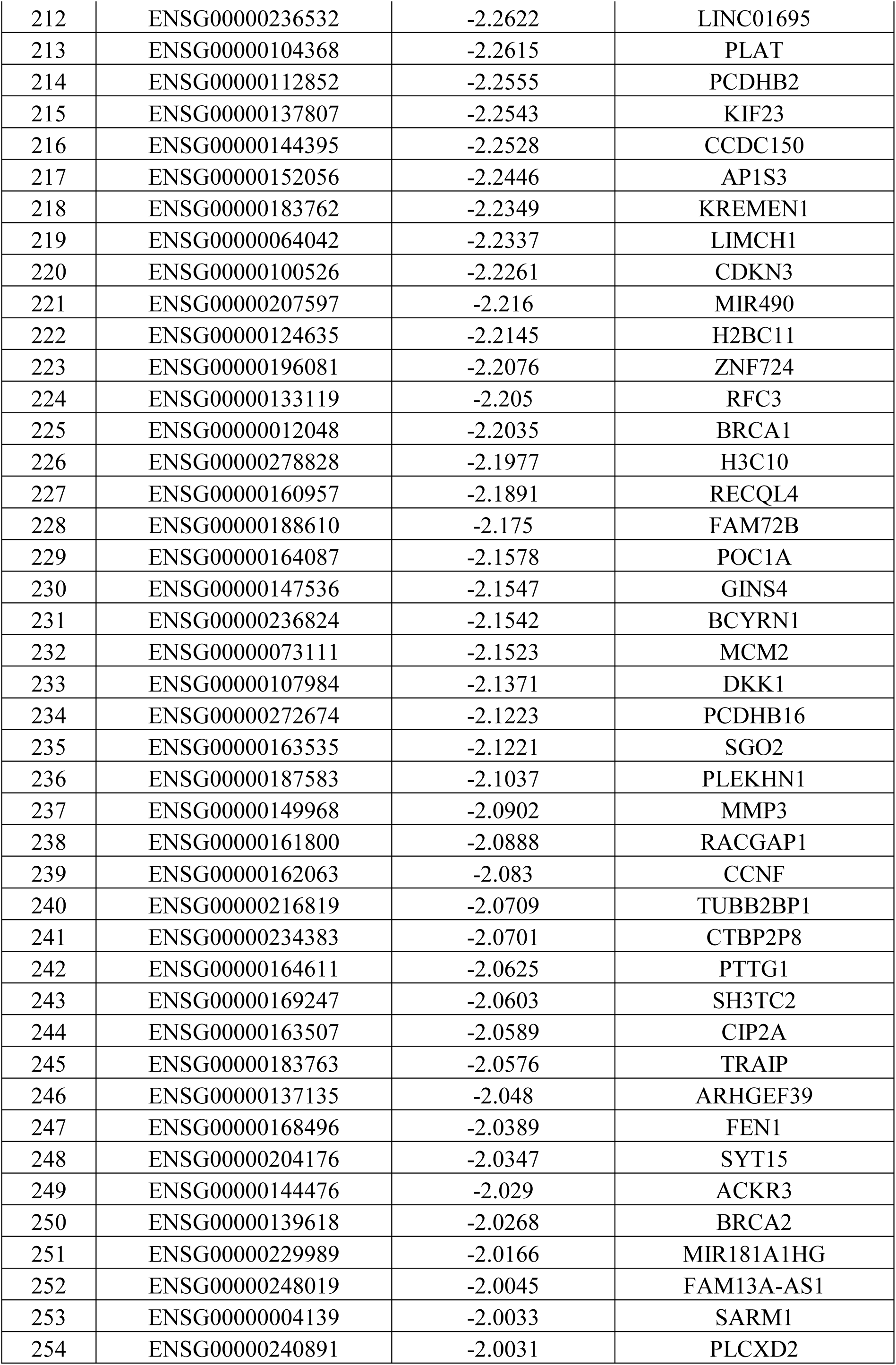
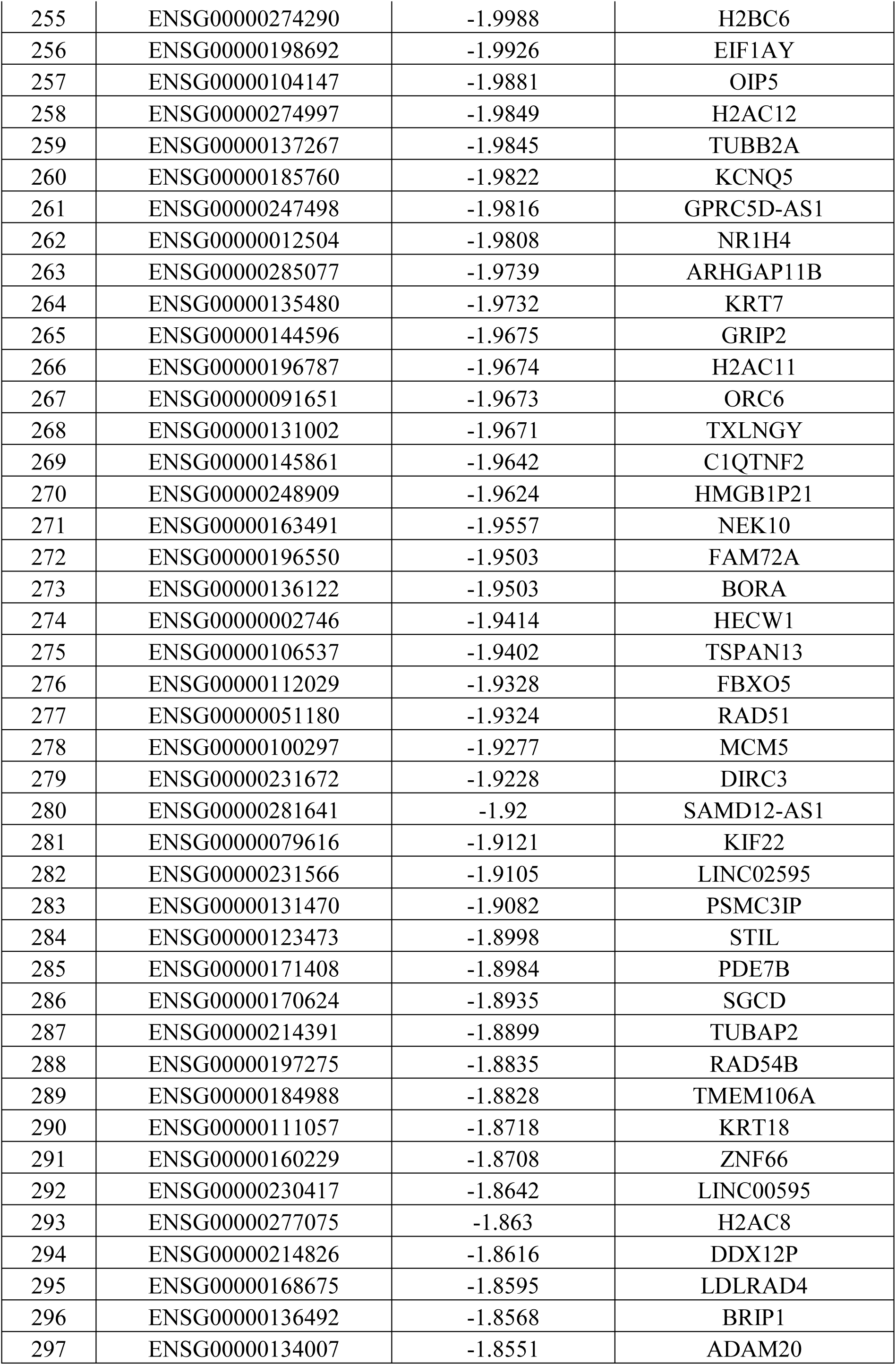
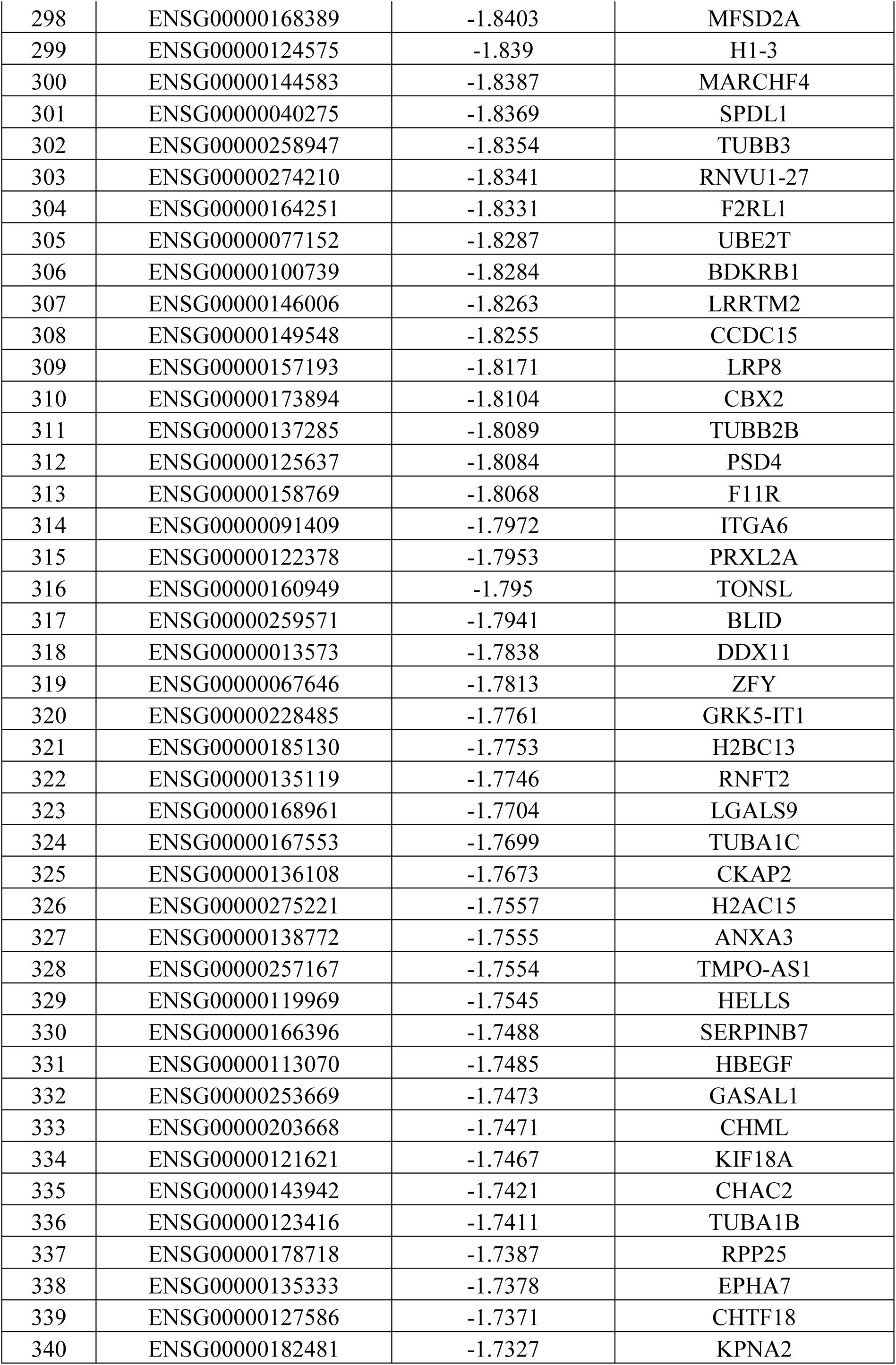
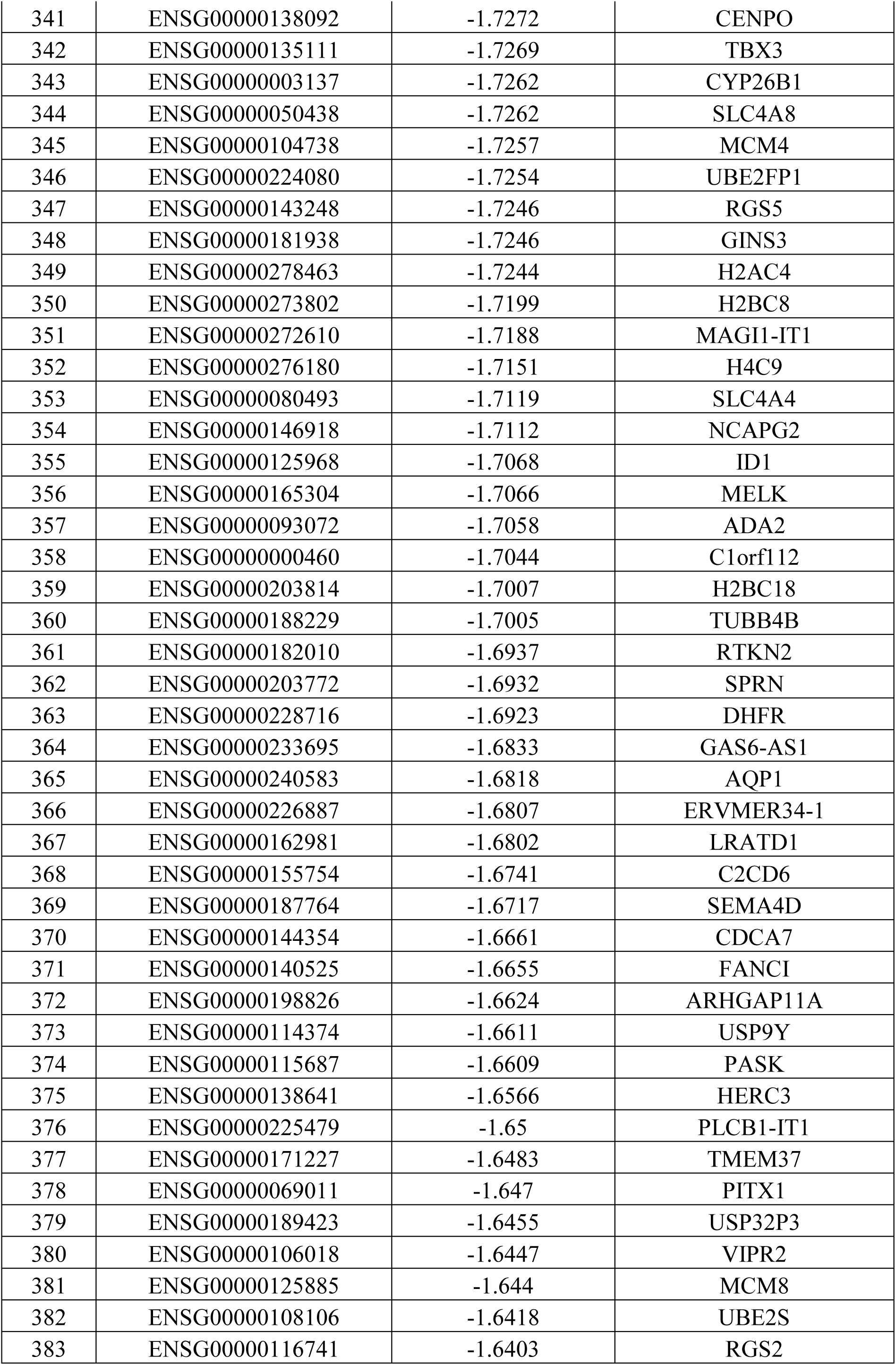
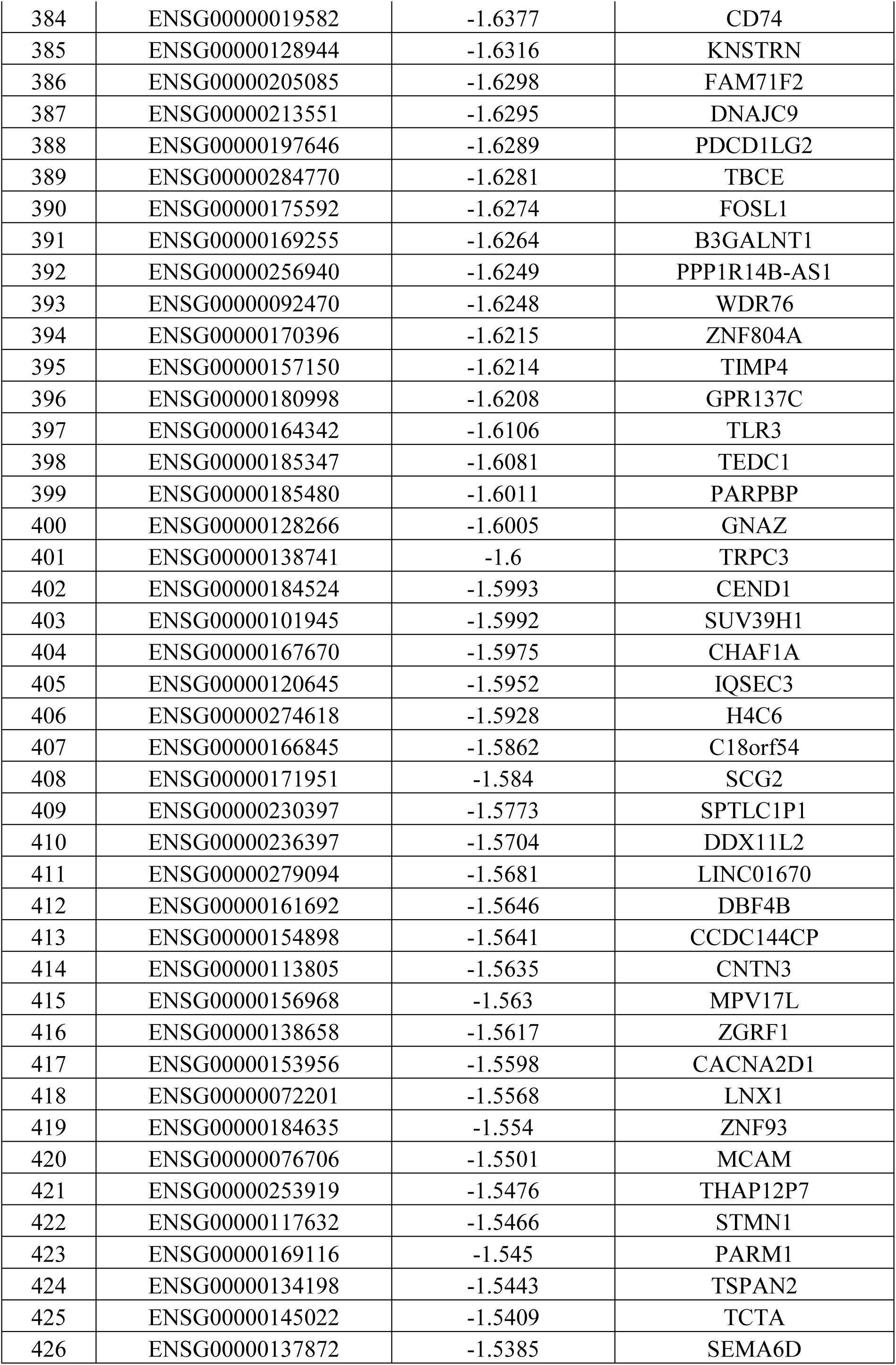
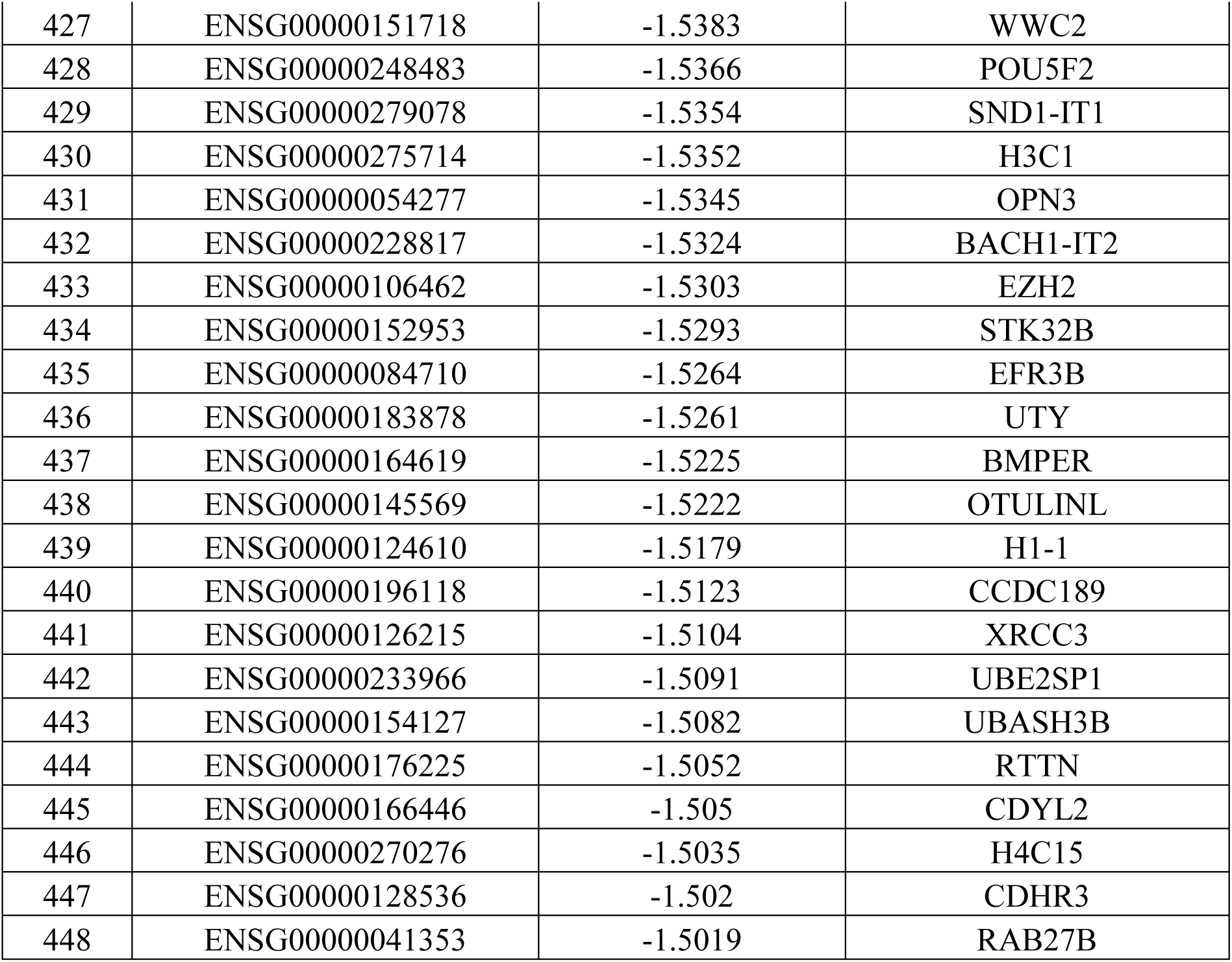
List of differentially expressed mRNAs and LINCRNAs in cancer associated fibroblasts compared to normal fibroblasts. The list shows the GeneID, relative fold change values (Log_2_FC) and gene symbol **1A:** List of mRNAs and LINCRNAs overexpressed in cancer associated fibroblasts as compared to normal fibroblasts (387) **1B:** List of mRNAs and LINCRNAs under-expressed in cancer associated fibroblasts as compared to normal fibroblasts (448)

### LINCRNA target prediction using NPInter V5.0

LincRNA targets were predicted for differentially expressed lincRNAs in normal compared to CAFs using NP Inter V5.0. This analysis yielded 31 miRNAs, 2 mRNA, 263 RNA-binding proteins (RBPs), and 1 ncRNA associated with a total of 14 lincRNAs Table 2. Also, there were 3 novel lincRNAs (LINC02344, LINC01670 and LINC02605) for which no data was found in the above database.

**Table 2:**
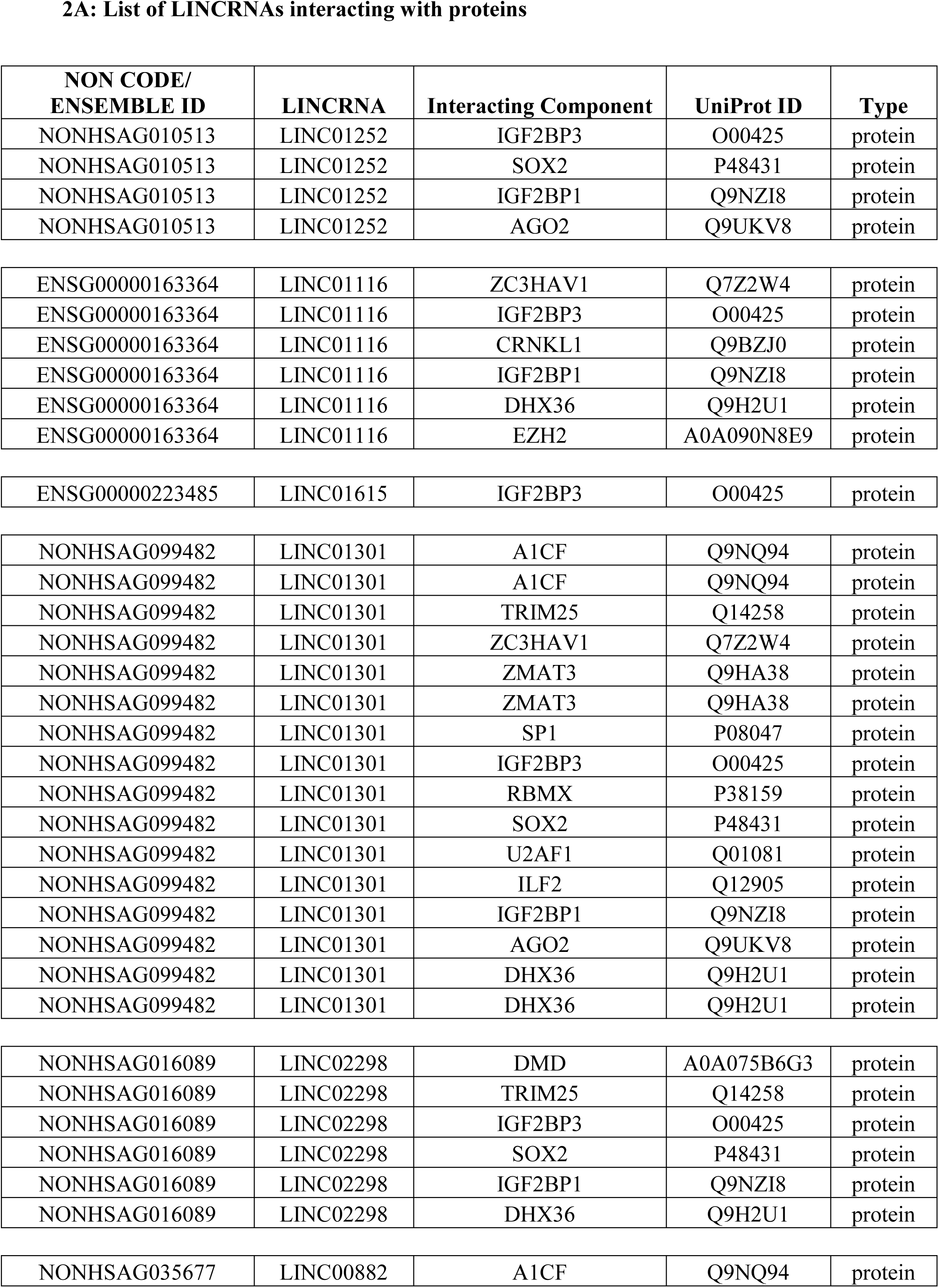
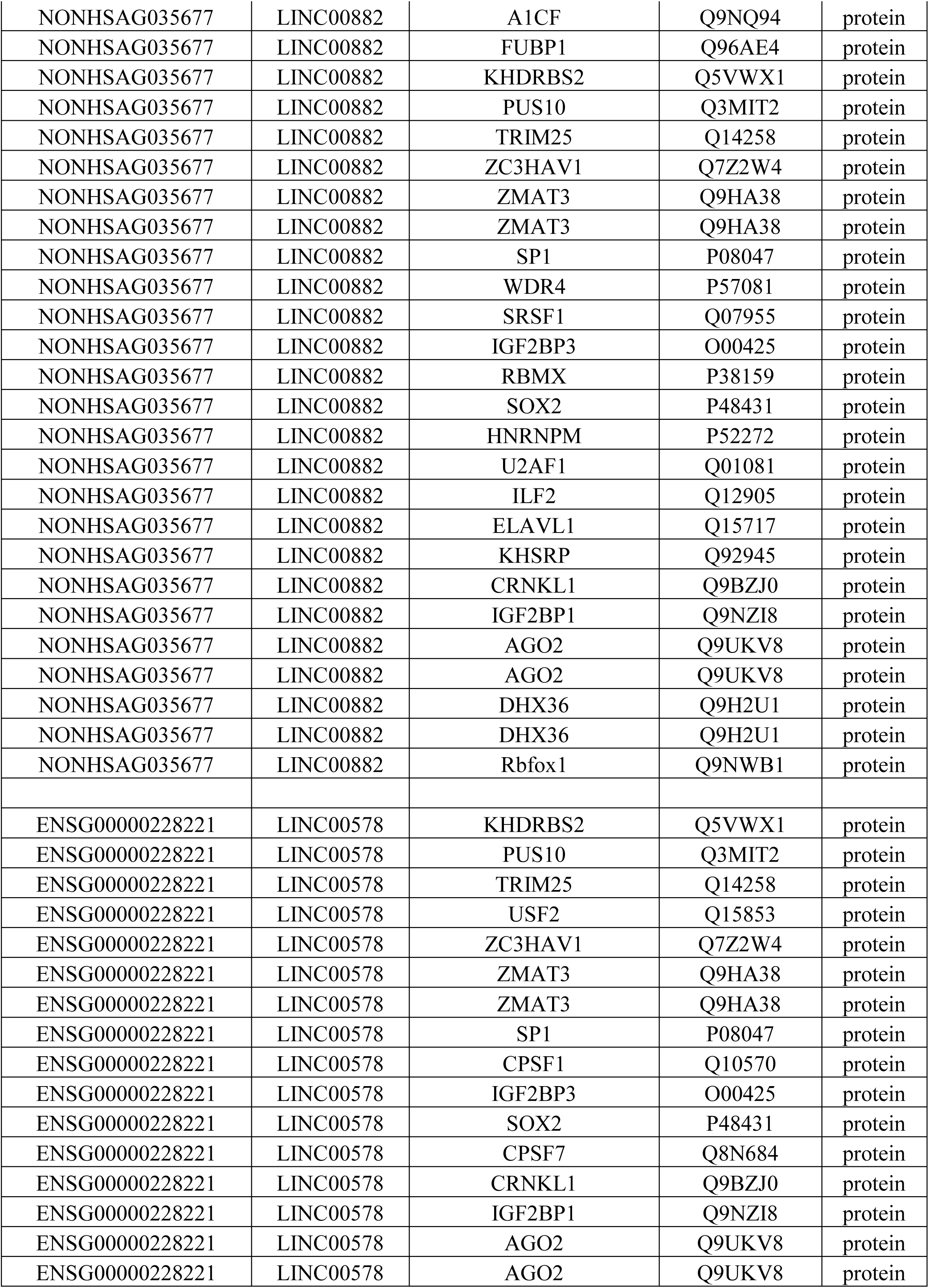
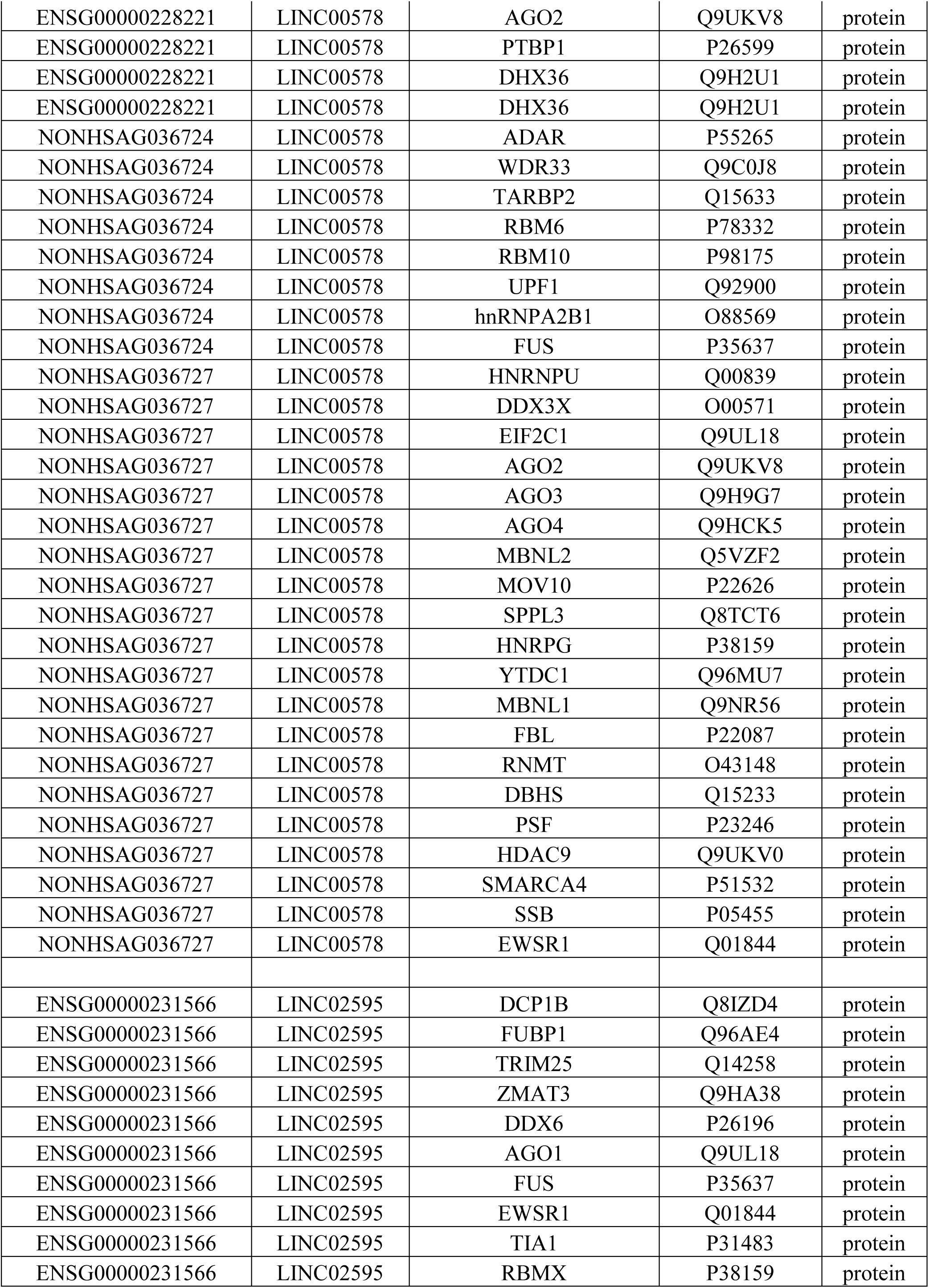
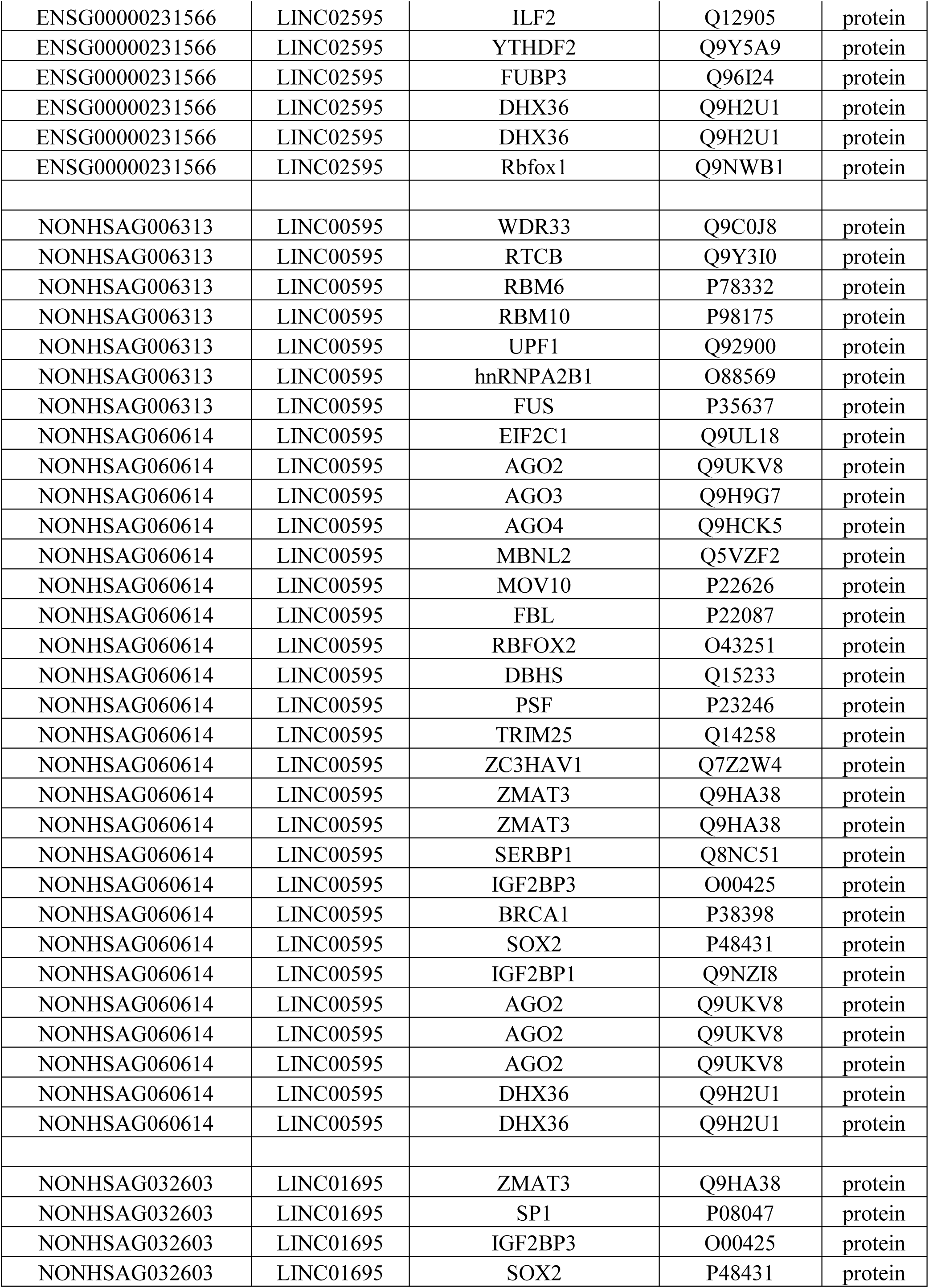
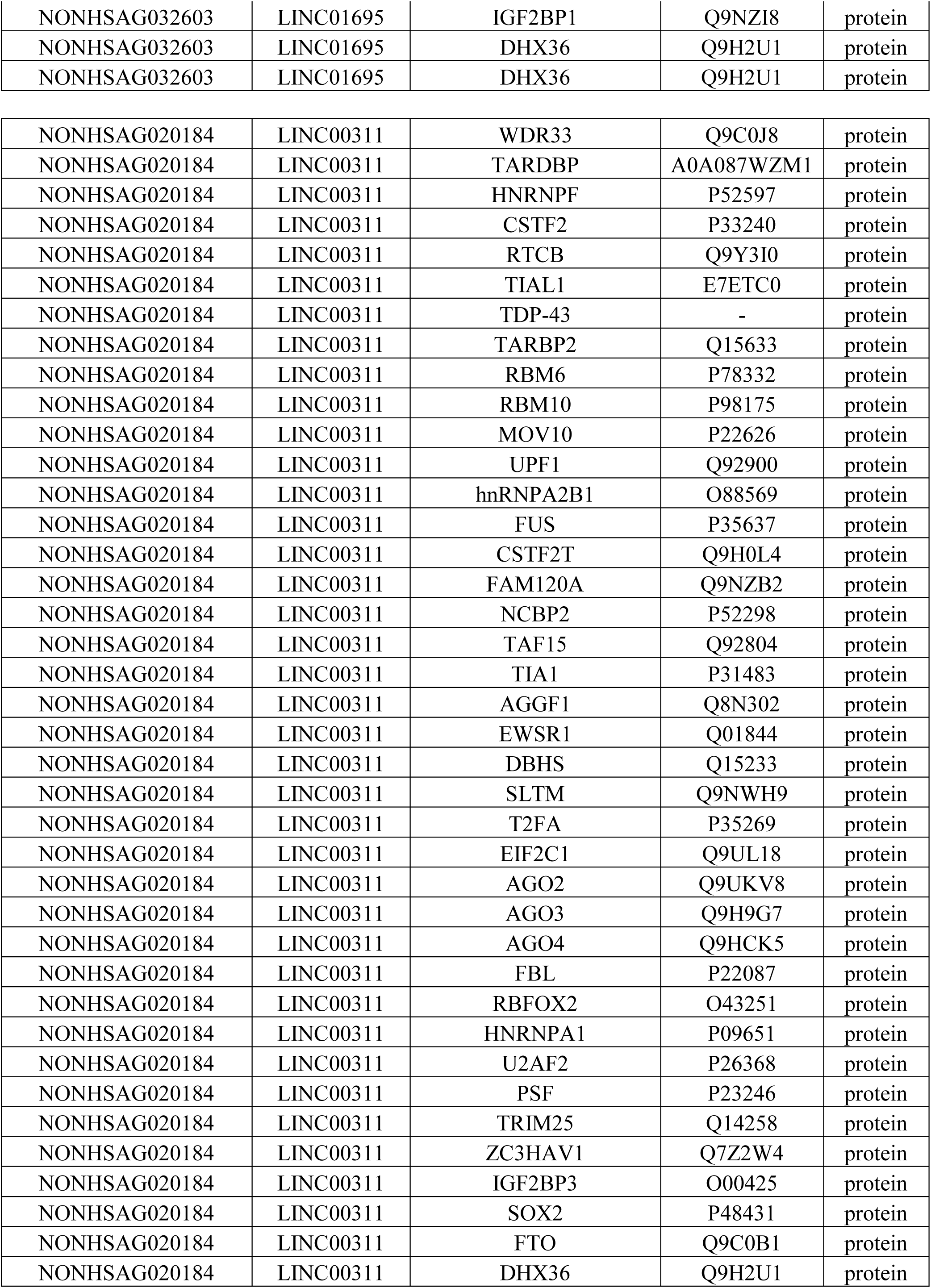
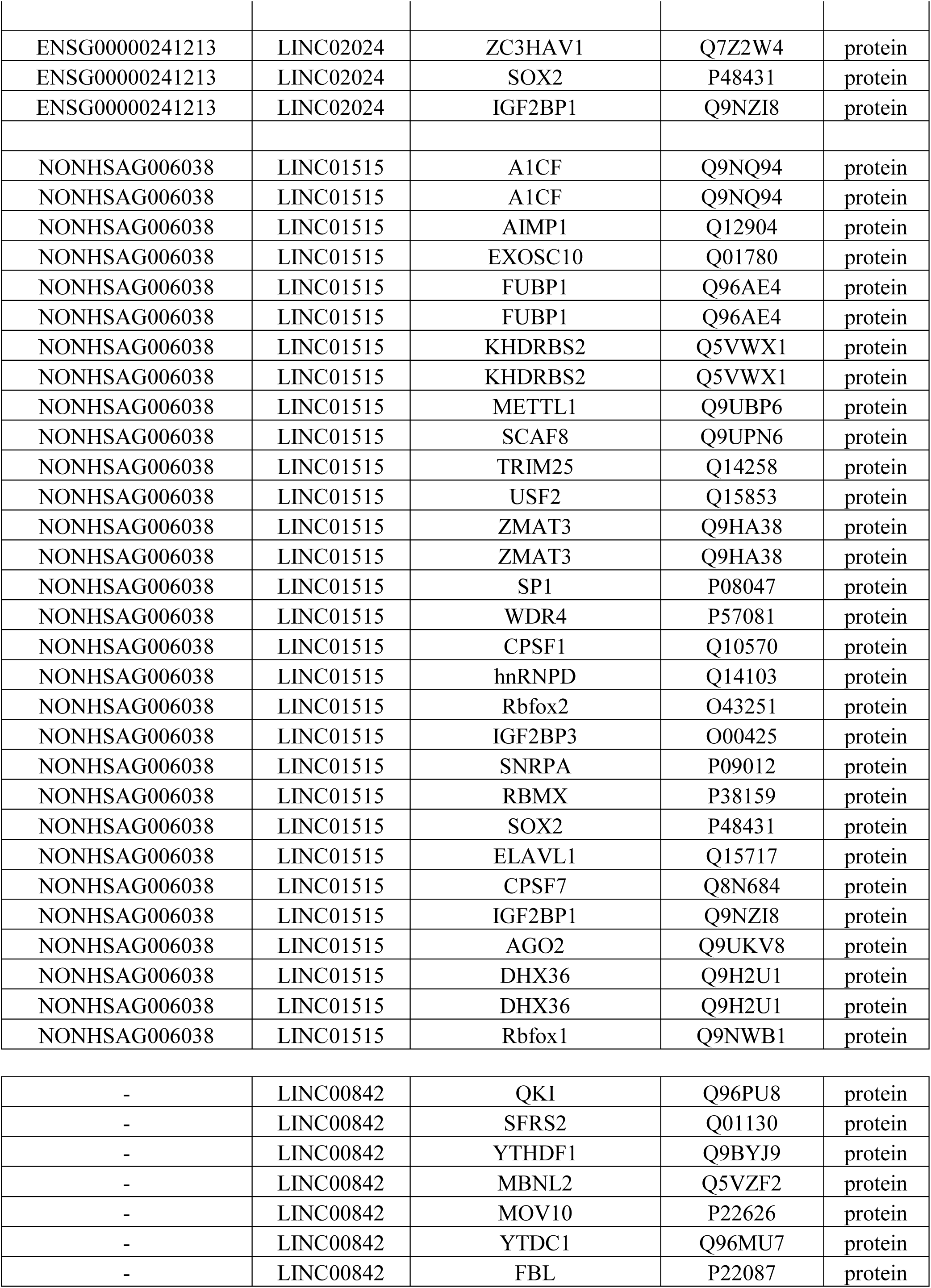
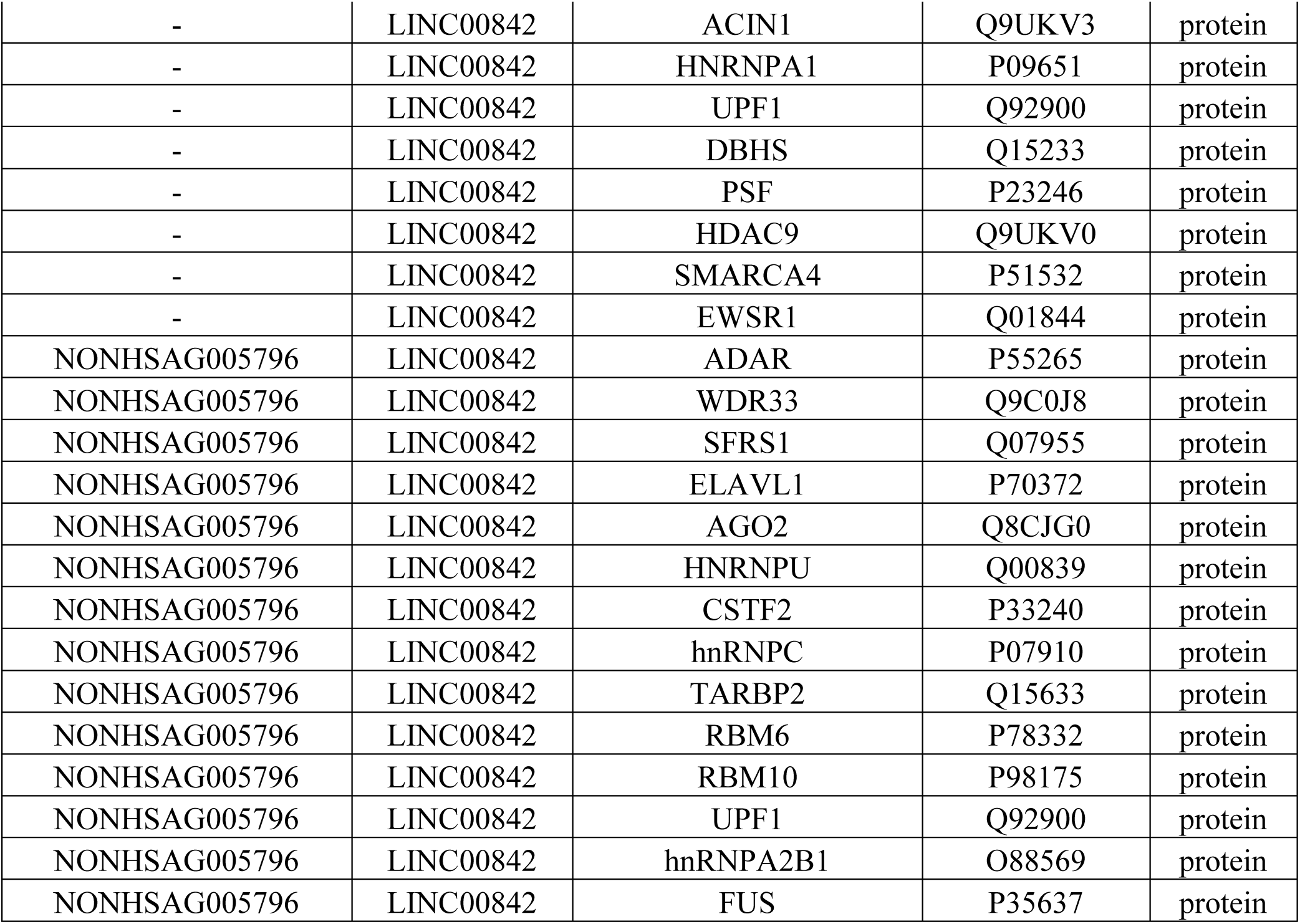
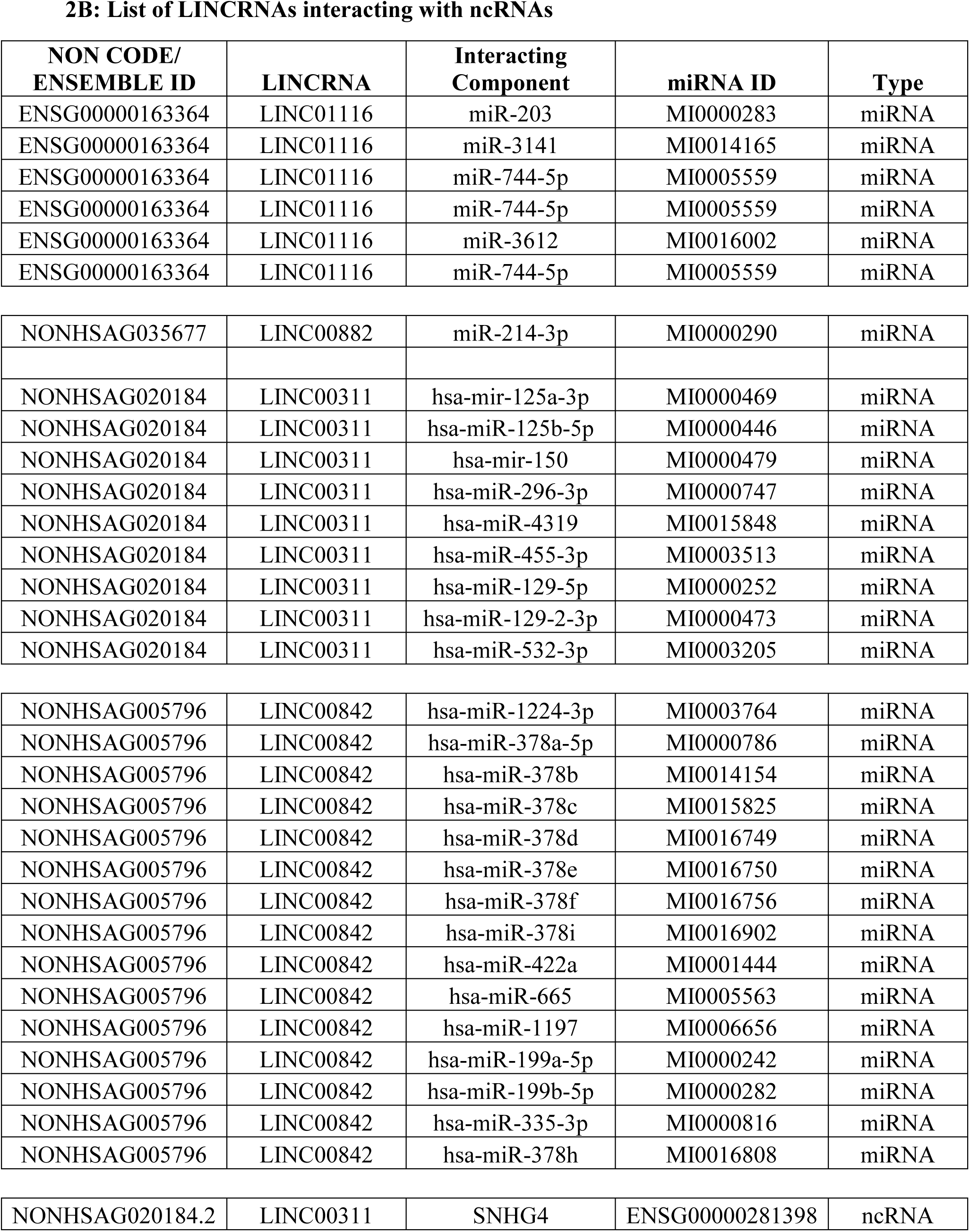
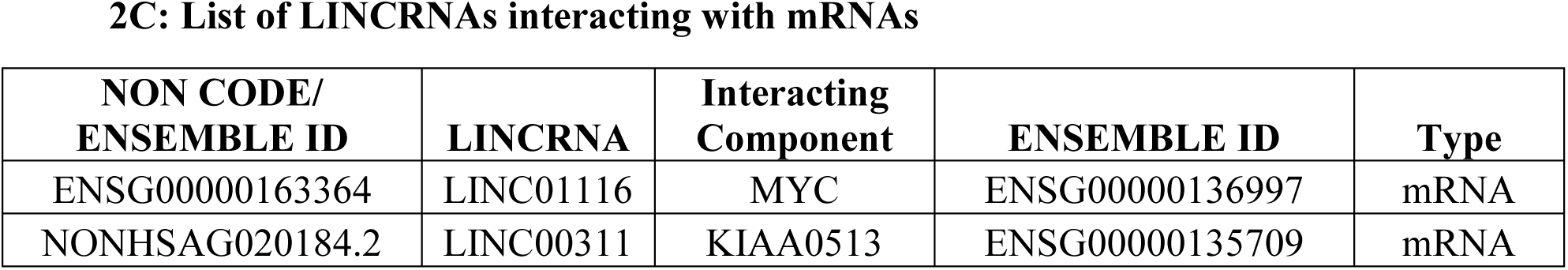
List of miRNAs, mRNAs and ncRNA associated with LINCRNAs. The list show Ensemble/NON CODE ID, interacting component and its Uniprot/miRNA ID and type of the interacting components **2A:** List of LINCRNAs interacting with proteins **2B:** List of LINCRNAs interacting with ncRNAs **2C:** List of LINCRNAs interacting with mRNA

### Prediction of targets of the miRNAs

The 24 miRNAs obtained from the above analysis targets were fed into miRDB for mRNA target prediction. mRNA targets with a score of >/= 95 were chosen. We have identified 288 mRNA targets associated with the input of 24 miRNAs.

The predicted mRNA targets were cross-referenced with our dataset to identify the common mRNAs. Subsequently, we established lincRNA-miRNA-mRNA combinations Table 3.

**Table 3:**
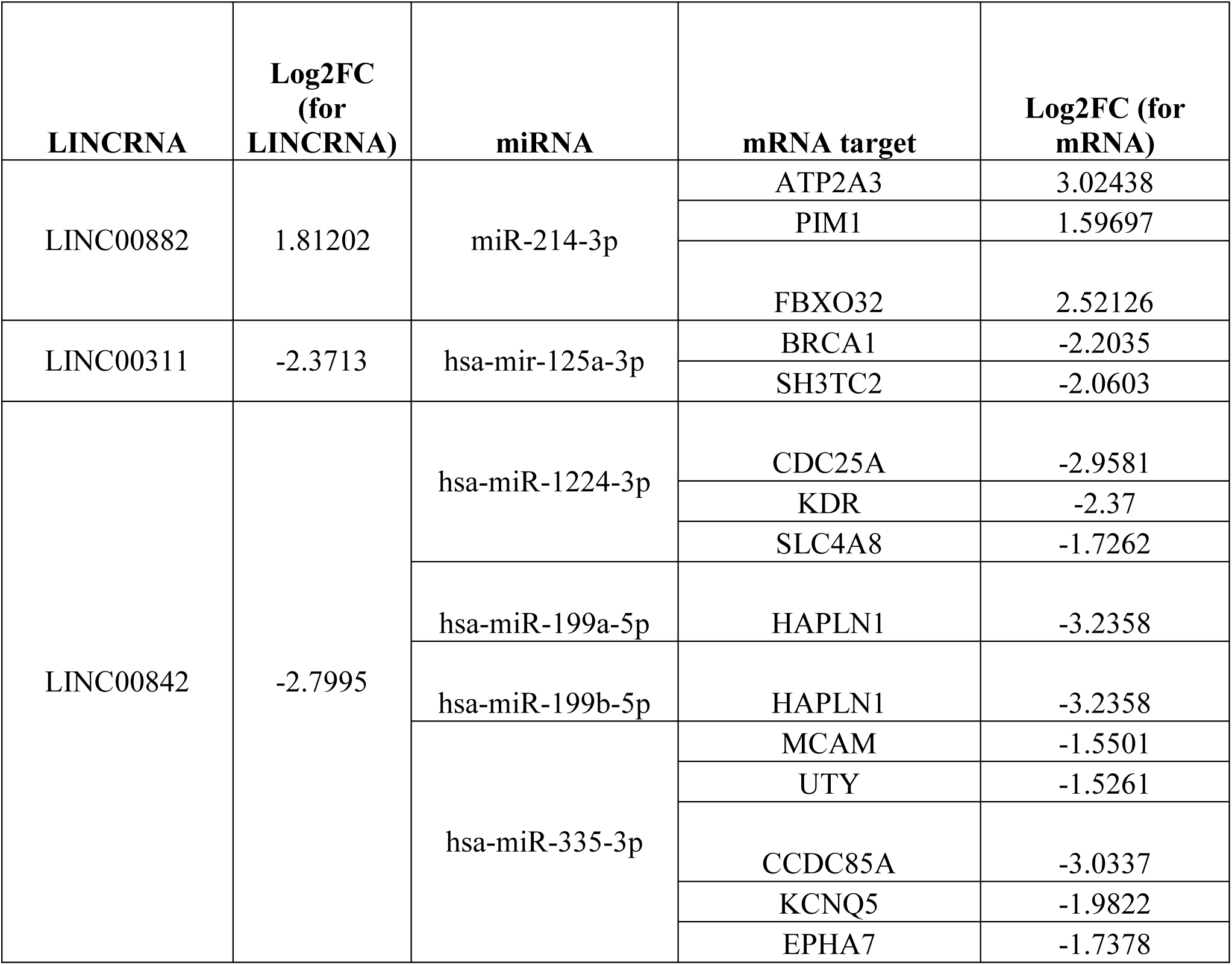
List of established lincRNA-miRNA-mRNA combinations. The list shows differentially expressed LINCRNA, relative fold change values of LINCRNA (Log_2_FC (for LINCRNA)), miRNA, mRNA target and relative fold change values of target mRNA (Log_2_FC (for mRNA)) in the same data set.

| **LINC RNA** | **Log2FC**<br>**(for**<br>**LINC RNA)** | **miRNA** | **mRNA target** | **Log2FC (for**<br>**mRNA)** |
| --- | --- | --- | --- | --- |
| LINC00882 | 1.81202 | miR-214-3p | ATP2A3 | 3.02438 |
|  |  |  | PIM1 | 1.59697 |
|  |  |  | FBXO32 | 2.52126 |
| LINC00311 | -2.3713 | hsa-mir-125a-3p | BRCA1 | -2.2035 |
|  |  |  | SH3TC2 | -2.0603 |
| LINC00842 | -2.7995 | hsa-miR-1224-3p | CDC25A | -2.9581 |
|  |  |  | KDR | -2.37 |
|  |  |  | SLC4A8 | -1.7262 |
|  |  | hsa-miR-199a-5p | HAPLN1 | -3.2358 |
|  |  | hsa-miR-199b-5p | HAPLN1 | -3.2358 |
|  |  | hsa-miR-335-3p | MCAM | -1.5501 |
|  |  |  | UTY | -1.5261 |
|  |  |  | CCDC85A | -3.0337 |
|  |  |  | KCNQ5 | -1.9822 |
|  |  |  | EPHA7 | -1.7378 |

## Discussion

The effect of microenvironment on the initiation, maintenance and progression of solid tumors has been established beyond doubt [14]. CAFs, which constitute a significant component of the TME are a major source of secreted factors. Interaction with the CAF and derived factors not only play a significant role in promoting tumorigenesis and metastasis but also influence the response of the tumor to drugs [15, 16]. It is likely that in response to chemotherapeutic drugs, some pro-tumorigenic actions of CAFs may be activated, which in turn aid the tumor cells in escaping from the drug challenge. Studies on prostate cancer have shown that tumor cells grown in the presence of CAFs or CAF-derived factors show much higher tolerance to drugs as compared to tumor cells grown alone[17]. Also, cells grown with CAF-derived factors have a higher potential to metastasize [18]. These highlight the possibility of targeting CAF/derived factors for therapeutic purposes. However, this requires the identification of specific factors and their effect on the tumor cells. CAFs are distinguished from normal fibroblasts by their contractile characteristics, and metabolic and transcriptomic activity [19, 20]. Also, they are shown to express higher levels of FAP, alpha SMA and vimentin [19, 21, 22]. However, till date, there are no known unique markers of CAF. Identification of such markers becomes extremely essential if CAFs/derived factors are to be targeted for therapy, particularly because there are both pro- and anti-tumor properties of these factors. In this study, we have used the NGS platform to do a comparative analysis of fibroblasts derived from non-malignant (BPH) and cancerous prostate. This study has identified 818 genes differentially expressed between normal and cancer-associated fibroblasts. Also, there are 17 lncRNAs which show differential expression.

Long Intergenic Non-Coding RNAs (lincRNAs) are RNA molecules exceeding 200 nucleotides, lacking protein-coding functions and non-overlapping with annotated coding genes. They impact gene expression by modulating chromatin structure, regulating transcription of nearby and distant genes, and interacting with DNA, RNA, and proteins [23–25] (Sup fig1). In cancer patients, differential lncRNA expression has been correlated with the overall survival (OS), metastasis, as well as tumor stage/grade [26–28]. LncRNAs have been detected in body fluids like plasma, serum, and urine using real-time PCR. One of the reasons lncRNAs are suitable as cancer diagnostic and prognostic biomarkers is their remarkable stability while circulating in body fluids, particularly when enclosed within exosomes or apoptotic bodies [29]. These characteristics of lncRNA make them attractive candidates for biomarkers. These biomarkers offer a minimally invasive alternative to conventional biopsies [30]. These markers can also be used to predict the prognosis of cancer patients, assess the risk of tumor metastasis and recurrence after surgery, and also to evaluate the success of therapeutic intervention. The distinct expression profiles of cancer-associated lncRNAs, which can vary significantly among different types of cancer, hold promise as efficient tumor biomarkers in various body fluids [27, 28, 31] (Supplementary table1 showing LncRNA as a prognostic and diagnostic marker and Supplementary Fig 4 showing tissue specific LNCRNA as potential biomarker). Despite the fact that lincRNAs are good biomarkers, targeting lincRNA or other ncRNA for therapeutic purposes have been extremely challenging. One of the reasons being very low conservation of lncRNAs across species. A small number of lncRNAs which are conserved between humans and mice have been discovered, while many human lncRNAs are absent in mice [32, 33].

Although it has been observed that lincRNAs show specific expression patterns in cancers, the heterogeneity in tumors makes it difficult to target them. Some studies have used in-silico approaches to identify lincRNA-miRNA-mRNA combinations. For example, a study has shown the influence of LOC101928304/miR-490-3p/LRRC2, a lincRNA-miRNA-mRNA axis on Atrial Fibrillation (AF). The levels of LOC101928304 and LRRC were elevated whereas miR-490-3p exhibited a decreased expression in the myocardial tissue of AF patients [34]. However, there is not much experimental data available. Given the advantages of using lincRNAs as biomarkers and also the difficulties in targeting them for therapeutic intervention, we feel that identifying a combination of lincRNA-miRNA-mRNA may provide better options for targeting. In our study we have predicted the targets of the differentially expressed lincRNAs and identified 15 lincRNA-miRNA-mRNA combinations. This would help in understanding the mechanism of action of these RNAs as well as identifying strategies for therapeutic targeting. However, this would in future need more experimental validation.

## Conflict of interest

Authors declare no conflict of interest

## Authors’ Contribution

AA analysed the transcriptomic data and prepared the manuscript draft; MSM, RRA and NN helped in collection of patient samples, deriving the fibroblasts and characterizing; VB and NT helped in collection of patient samples and clinical evaluation; RK helped with all the patient related work and helped in procuring funding; PR conceived and strategized the study and procured the funding.

## Financial Support

This study was supported by Indian Council for Medical Research, Govt of India (2019-0937). AA is supported by Lady Tata Memorial Trust Fellowship. The authors thank Centre for Human Genetics, Bengaluru and Institute of Nephro-Urology, Bengaluru for all the support during the course of this study.

## Supplementary material with legends

**Sup table 1:** LNCRNAs as cancer prognostic and/or diagnostic marker. The table shows the type of cancer, LNCRNA involved, LNCRNA expression, their relevance as prognostic or diagnostic marker.

**Sup fig 1:** LNCRNA mediated gene expression regulation

**Sup fig 2:** RNA sequencing analysis workflow

**Sup fig 3:** LINCRNA analysis work flow

**Sup fig 4:** Tissue specific LNCRNA as potential biomarkers

## Notes

### Competing Interest Statement

The authors have declared no competing interest.

## References

1. Patel, H., et al., Modulating secreted components of tumor microenvironment: A masterstroke in tumor therapeutics. Cancer Biol Ther, 2018. 19(1): p. 3–12.

2. Baghban, R., et al., Tumor microenvironment complexity and therapeutic implications at a glance. Cell Commun Signal, 2020. 18(1): p. 59.

3. Anderson, N.M. and M.C. Simon, The tumor microenvironment. Curr Biol, 2020. 30(16): p. R921–R925.

4. Xiao, Y. and D. Yu, Tumor microenvironment as a therapeutic target in cancer. Pharmacol Ther, 2021. 221: p. 107753.

5. Brennen, W.N., J.T. Isaacs, and S.R. Denmeade, Rationale behind targeting fibroblast activation protein-expressing carcinoma-associated fibroblasts as a novel chemotherapeutic strategy. Mol Cancer Ther, 2012. 11(2): p. 257–66.

6. Mao, X., et al., Crosstalk between cancer-associated fibroblasts and immune cells in the tumor microenvironment: new findings and future perspectives. Mol Cancer, 2021. 20(1): p. 131.

7. Bu, L., et al., Functional diversity of cancer-associated fibroblasts in modulating drug resistance. Cancer Sci, 2020. 111(10): p. 3468–3477.

8. Asif, P.J., et al., The Role of Cancer-Associated Fibroblasts in Cancer Invasion and Metastasis. Cancers (Basel), 2021. 13(18).

9. Jena, B.C., et al., Cancer associated fibroblast mediated chemoresistance: A paradigm shift in understanding the mechanism of tumor progression. Biochim Biophys Acta Rev Cancer, 2020. 1874(2): p. 188416.

10. Rizzolio, S., S. Giordano, and S. Corso, The importance of being CAFs (in cancer resistance to targeted therapies). J Exp Clin Cancer Res, 2022. 41(1): p. 319.

11. Zheng, Y., et al., NPInter v5.0: ncRNA interaction database in a new era. Nucleic Acids Res, 2023. 51(D1): p. D232–D239.

12. NPInter. Available from: http://bigdata.ibp.ac.cn/npinter5/.

13. miRDB. Available from: https://mirdb.org/.

14. Joyce, J.A. and J.W. Pollard, Microenvironmental regulation of metastasis. Nat Rev Cancer, 2009. 9(4): p. 239–52.

15. Sun, Y., et al., SFRP2 augments WNT16B signaling to promote therapeutic resistance in the damaged tumor microenvironment. Oncogene, 2016. 35(33): p. 4321–34.

16. Qiao, Y., et al., IL6 derived from cancer-associated fibroblasts promotes chemoresistance via CXCR7 in esophageal squamous cell carcinoma. Oncogene, 2018. 37(7): p. 873–883.

17. Cheteh, E.H., et al., Human cancer-associated fibroblasts enhance glutathione levels and antagonize drug-induced prostate cancer cell death. Cell Death Dis, 2017. 8(6): p. e2848.

18. Linxweiler, J., et al., Cancer-associated fibroblasts stimulate primary tumor growth and metastatic spread in an orthotopic prostate cancer xenograft model. Sci Rep, 2020. 10(1): p. 12575.

19. Xing, F., J. Saidou, and K. Watabe, Cancer associated fibroblasts (CAFs) in tumor microenvironment. Front Biosci (Landmark Ed), 2010. 15(1): p. 166–79.

20. De Wever, O., et al., Stromal myofibroblasts are drivers of invasive cancer growth. Int J Cancer, 2008. 123(10): p. 2229–38.

21. Gilardi, L., et al., Imaging Cancer-Associated Fibroblasts (CAFs) with FAPi PET. Biomedicines, 2022. 10(3).

22. Muchlińska, A., et al., Alpha-smooth muscle actin-positive cancer-associated fibroblasts secreting osteopontin promote growth of luminal breast cancer. Cell Mol Biol Lett, 2022. 27(1): p. 45.

23. Ti, W., J. Wang, and Y. Cheng, The Interaction Between Long Non-Coding RNAs and Cancer-Associated Fibroblasts in Lung Cancer. Front Cell Dev Biol, 2021. 9: p. 714125.

24. Ransohoff, J.D., Y. Wei, and P.A. Khavari, The functions and unique features of long intergenic non-coding RNA. Nat Rev Mol Cell Biol, 2018. 19(3): p. 143–157.

25. Sebastian-delaCruz, M., et al., The Role of lncRNAs in Gene Expression Regulation through mRNA Stabilization. Noncoding RNA, 2021. 7(1).

26. Qian, Y., L. Shi, and Z. Luo, Long Non-coding RNAs in Cancer: Implications for Diagnosis, Prognosis, and Therapy. Front Med (Lausanne), 2020. 7: p. 612393.

27. Gao, N., et al., Long Non-Coding RNAs: The Regulatory Mechanisms, Research Strategies, and Future Directions in Cancers. Front Oncol, 2020. 10: p. 598817.

28. Beylerli, O., et al., Long noncoding RNAs as promising biomarkers in cancer. Noncoding RNA Res, 2022. 7(2): p. 66–70.

29. Akers, J.C., et al., Biogenesis of extracellular vesicles (EV): exosomes, microvesicles, retrovirus-like vesicles, and apoptotic bodies. J Neurooncol, 2013. 113(1): p. 1–11.

30. Su, Y.J., et al., Circulating Long Noncoding RNA as a Potential Target for Prostate Cancer. Int J Mol Sci, 2015. 16(6): p. 13322–38.

31. Bolha, L., M. Ravnik-Glavač, and D. Glavač, Long Noncoding RNAs as Biomarkers in Cancer. Dis Markers, 2017. 2017: p. 7243968.

32. Necsulea, A., et al., The evolution of lncRNA repertoires and expression patterns in tetrapods. Nature, 2014. 505(7485): p. 635–40.

33. Lee, J.T., Epigenetic regulation by long noncoding RNAs. Science, 2012. 338(6113): p. 1435–9.

34. Ke, X., et al., Construction and Analysis of the lncRNA-miRNA-mRNA Network Based on Competing Endogenous RNA in Atrial Fibrillation. Front Cardiovasc Med, 2022. 9: p. 791156.

