## Supplementary figures and images for "Comparative transcriptome of normal and cancer-associated fibroblasts"

### Supplementary Figure 1

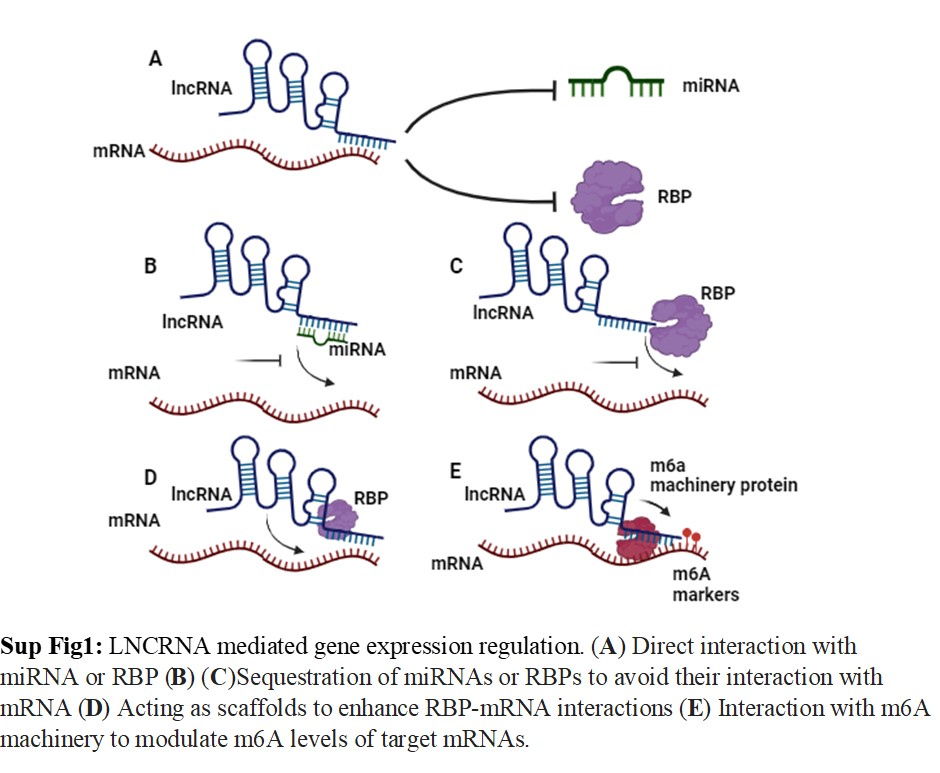

### Supplementary Figure 2

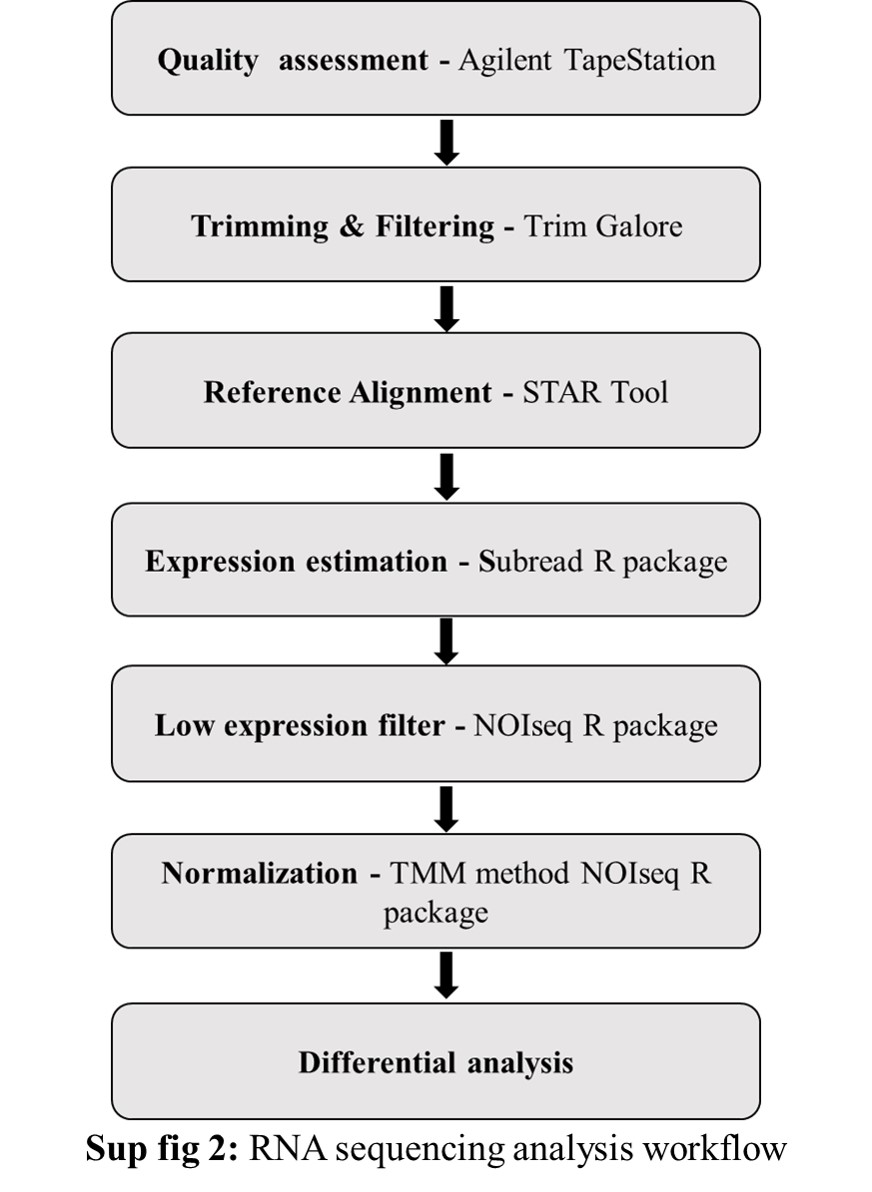

### Supplementary Figure 3

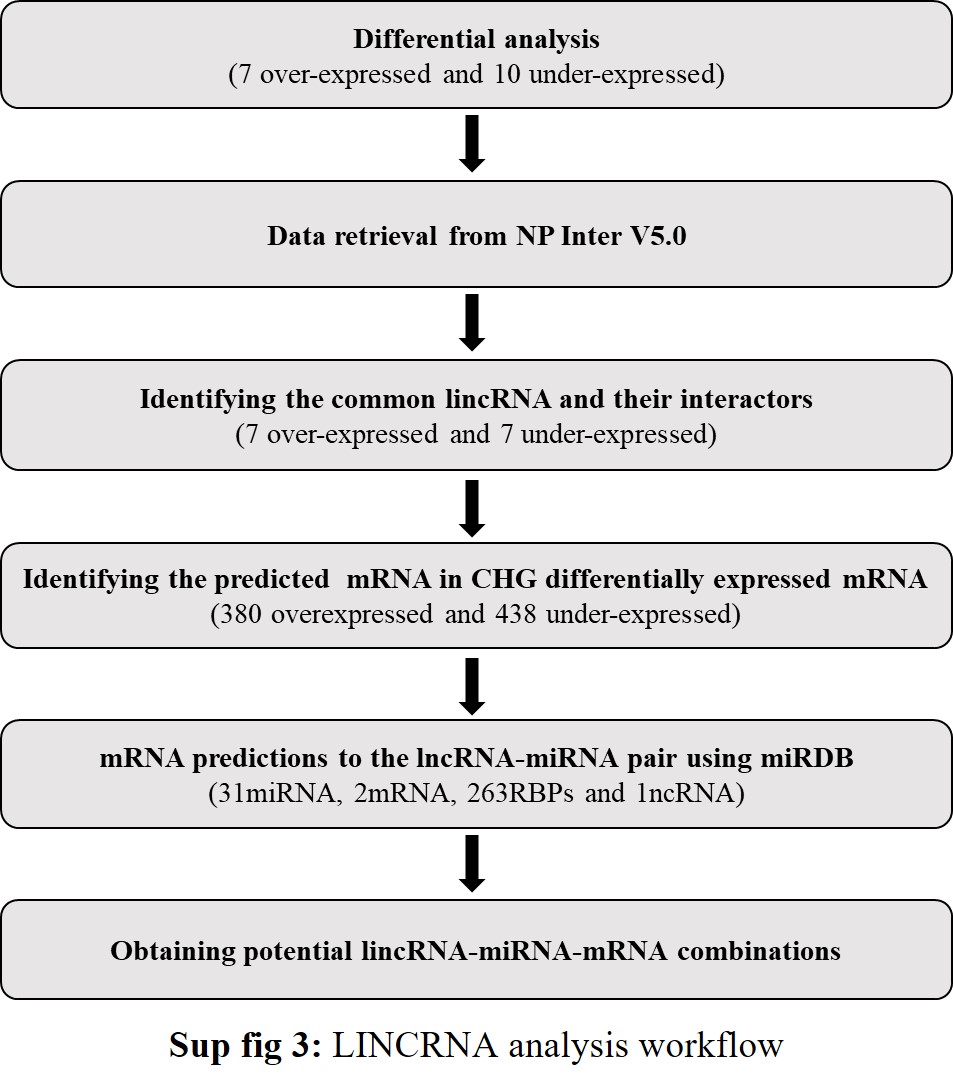

### Supplementary Figure 4

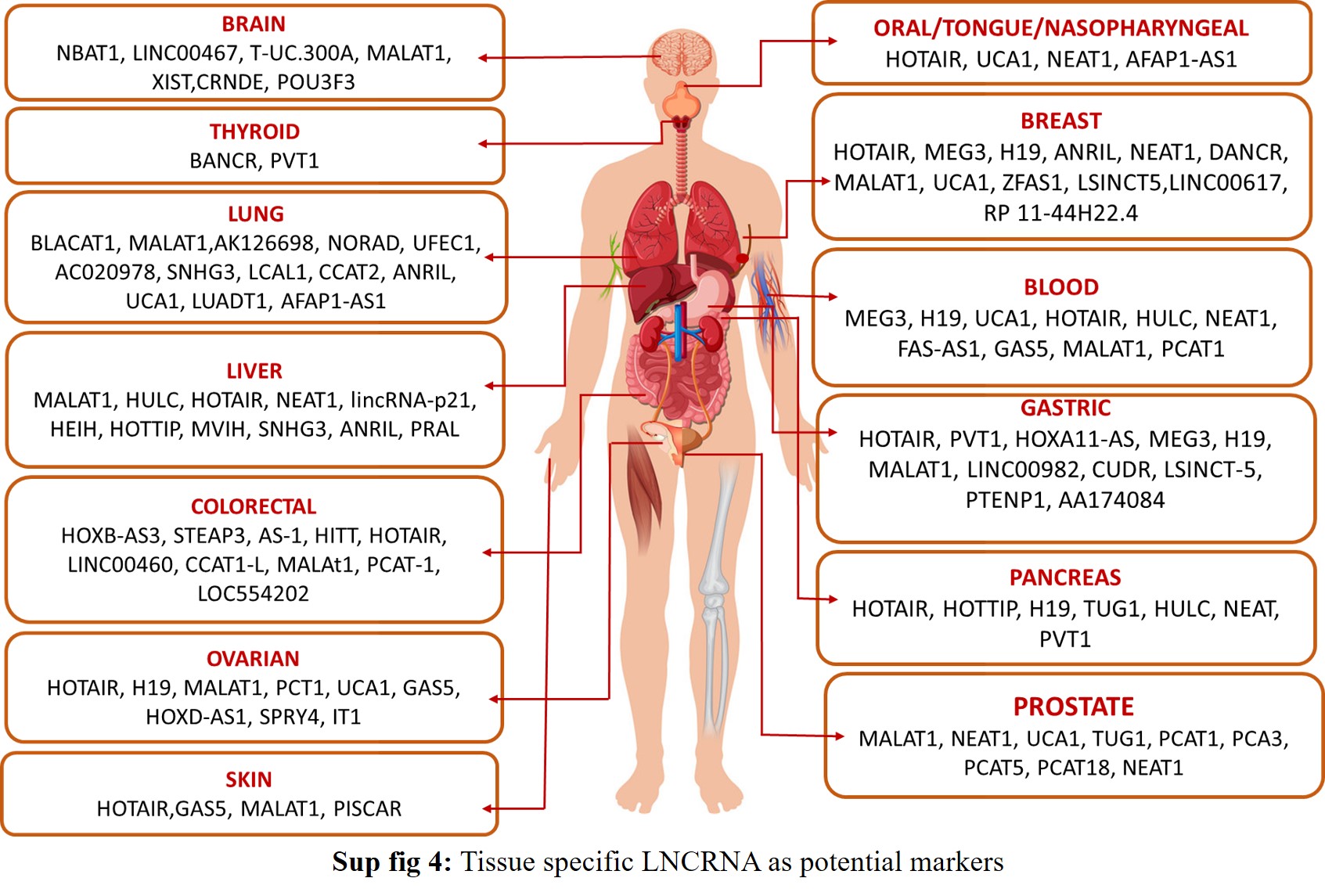
