## Supplementary Table 1 for "Comparative transcriptome of normal and cancer-associated fibroblasts"

| Sup table 1: LNCRNAs as cancer prognostic and/or diagnostic marker | | | | |
|---|---|---|---|---|
| **Cancer Type** | **LNCRNA** | **over/under expressed** | **Diagnosis/Prognosis** | **Reference** |
| Prostate | PCA3 | Over | Diagnosis | PMID: 31189930,PMID: 26080435 |
| | MALAT1 | Over | Diagnosis | PMID: 23726266 |
| | PVT1 | Over | Prognosis | PMID: 31766781 |
| | GAS5-007 | Over | Prognosis, Therapeutic Target | PMID: 28771526 |
| | ScHLAP1 | Over | Prognosis | PMID: 34107378 |
| | LincRNA-p21 | Over | Diagnosis | PMID: 31189930 |
| Bladder | PTENP1 | Under | Diagnosis | PMID: 30285771 |
| | UCA1 | Over | Diagnosis | PMID: 26544536 |
| | SPRY4-IT1 | Over | Diagnosis | PMID: 27998761 |
| | HOTAIR | Over | Prognosis | PMID: 26469956 |
| Breast | H19 | Over | Diagnosis | PMID: 27540977 |
| Colorectal | CCAT1 | Over | Diagnosis | PMID: 24777251 |
| | CCAT2 | Over | Prognosis | PMID: 23796952 |
| | MALAT1 | Over | Prognosis | PMID: 21503572 |
| | MEG3 | Under | Prognosis | PMID: 14602737 |
| | HOTAIR | Over | Prognosis | PMID: 21862635 |
| Gastric | HOTAIR | Over | Diagnosis | PMID: 32951010 |
| | HOTAIR | Over | Prognosis | PMID: 25280565 |
| | MALAT1 | Over | Diagnosis | PMID: 28942451 |
| | MALAT1 | Over | Prognosis | PMID: 27486823 |
| Liver | MALAT1 | Over | Prognosis | PMID: 30564069 |
| | H19 | Over | Diagnosis | PMID: 27540977 |
| Esophageal | CCAT2 | Over | Prognosis | PMID: 27540977 |
| | PCAT1 | Over | Prognosis | PMID: 25731728 |
| Glioma | CASC2 | Under | Diagnosis | PMID: 25919911 |
| | CRNDE | Over | Prognosis | PMID: 25813405 |
| Thyroid | HOTAIR | Over | Diagnosis | PMID: 28670492 |
